# Genetic ancestry in Puerto Rican Afro-descendants illustrates diverse histories of African diasporic populations

**DOI:** 10.1101/2024.09.02.610898

**Authors:** Maria A. Nieves-Colón, Emma C. Ulrich, Lijuan Chen, Gabriel A. Torres Colón, Maricruz Rivera Clemente, La Corporación Piñones Se Integra (COPI), Jada Benn Torres

**Affiliations:** Department of Anthropology, University of Minnesota Twin Cities, Minneapolis, MN 55455; Department of Anthropology, Genetic Anthropology and Biocultural Studies Laboratory, Vanderbilt University, Nashville, TN 37235; La Corporación Piñones Se Integra (COPI) Loíza, Puerto Rico; Vanderbilt Genetics Institute, Vanderbilt University Nashville, TN 37235

**Author notes:** These authors contributed equally (Maria A. Nieves-Colón, Emma C. Ulrich).

**Keywords:** genomics, African ancestry, Puerto Rico, African diaspora

## Abstract

**Objectives:** Genetic studies of contemporary Puerto Ricans reflect a demographic history characterized by admixture between Indigenous American, African, and European peoples. While previous studies provide genetic perspectives on the general Puerto Rican population, less is known about the island’s sub-populations, specifically Afro-Puerto Ricans.

**Materials and Methods:** In this study, the genetic ancestry of Afro-Puerto Ricans is characterized and compared to other Caribbean populations. Thirty DNA samples collected among self-identified Puerto Ricans of African descent in Loíza (n=2), Piñones (n=13), San Juan (n=2), Mayagüez (n=9), and Ponce (n=4), were genotyped at 750,000 loci on the National Geographic Genochip. We then applied unsupervised clustering and dimensionality-reduction methods to detect continental and subcontinental African and European genetic ancestry patterns.

**Results:** Admixture analyses reveal that on average, the largest genetic ancestry component for Afro-Puerto Ricans is African in origin, followed by European and Indigenous American genetic ancestry components. African biogeographic origins of Afro-Puerto Ricans align most closely with contemporary peoples of Lower Guinea and the Bight of Biafra, while the European genetic ancestry component is most similar to contemporary Iberian, Italian, and Basque populations. These findings contrast with the biogeographic origins of comparative Barbadian and Puerto Rican populations.

**Discussion:** Our results suggest that while there are similarities in general patterns of genetic ancestry among African descendants in the Caribbean, there is previously unrecognized regional heterogeneity, including among Puerto Rican sub-populations. These results are also consistent with available historical sources, while providing depth absent from the documentary record, particularly with regard to African ancestry.

## 1. Introduction

Located where the Greater and Lesser Antilles meet in the Caribbean Sea, Puerto Rico, an island of approximately 9065 sq km (3500 sq miles), is home to an estimated 3 million people (Central Intelligence, 2021). Since initially inhabited approximately 5-6000 years ago (Pestle, Perez, & Koski-Karell, 2023; Rodríguez Ramos, Rodríguez López, & Pestle, 2023), the island has continuously been populated and has served as a cross-roads where Indigenous Caribbean populations interacted with each other and then later beginning in the late 15^th^ century with migrants from Africa, Asia, and Europe.

Human presence on Puerto Rico has been documented via archeological analyses (Curet, 2005; Siegel, 2005), and more recently using genomic perspectives (Fernandes et al., 2021; Nägele et al., 2020; Nieves-Colón et al., 2020). The overall picture provided by these and related studies indicate that the Caribbean region was initially populated through serial migratory pulses from South, and possibly Central, America. As a result of these dispersion events, Native populations settled the Antilles and developed complex social, political, and economic systems (Hofman et al., 2020; Laffoon et al., 2014; Siegel, 2005). The arrival of European colonial settlers at the end of the 15^th^ century severely impacted, but did not eliminate, Indigenous Caribbean populations.

Characteristic of Spanish colonization in the Americas, economic interests often dictated how the Indigenous populations and island resources were exploited. Forced labor systems that exploited Indigenous and enslaved African individuals were essential to the development of Puerto Rican colonial commercial industries. In the Spanish Caribbean, trade in enslaved Africans was legalized in 1501, less than a decade after the European discovery of the Americas. Due to the declining Indigenous populations available for exploitative labor, enslaved Africans were forcibly migrated into Puerto Rico starting around 1511 to work within burgeoning mining industries, pearl fisheries, and agricultural sectors (Stark, 2009; Wheat, 2016). The use of enslaved African labor was also precipitated by shifts in colonists’ sentiments regarding the suitability of African over Indigenous laborers (Las Casas, 2004). Advocacy of enslaved African labor was endorsed by the Catholic church, as Dominican missionaries owned and used enslaved peoples to work church-owned farms and ranches throughout the 19^th^ century (García Leduc, 2002). Indeed, within the first three decades of the 16^th^ century, city officials in San Juan, Puerto Rico acknowledged the significant role that enslaved African labor had in colonial expansion and settlement, citing the use of enslaved peoples as a “necessary evil” (Wheat, 2016, p. 5).

Institutionalized slavery in Puerto Rico lasted throughout the late 19th century, was outlawed in 1873, and ended in practice after a three year apprenticeship period on April 20, 1876 (Rodríguez-Silva, 2012). Over the course of legalized slavery, enslaved Africans and African descendants were never a demographic majority, numbering in the thousands at any given point. As indicated in archival records, from 1776 until abolition, the highest concentrations of enslaved persons were consistently distributed between three municipalities: Mayagüez, Ponce, and Guayama (Figueroa, 2005). The geographic origins of enslaved Africans that were trafficked to Puerto Rico varied over time and by the enslaving colonial power. For example, throughout the 16^th^ century, Spanish slavers took people from Greater Senegambia then later, during the 17^th^ century, from West Central Africa and Angola. In the early 1700’s, French slavers seized control of the trade and imported Africans from Upper Guinea into Puerto Rico. Prior to the mid-18^th^ century, the British gained control of the trade and trafficked people from the Bight of Biafra (Stark, 2009; Wheat, 2016). The changing origins of enslaved African persons in Puerto Rico is also documented in the Transatlantic Slave Trade database (https://www.slavevoyages.org/voyage/database). Per this database, from 1761-1850 approximately 13,308 Africans disembarked in Puerto Rico coming from either Sierra Leone or further south along the western coast in the Bight of Biafra (Eltis, 2009). Archival research suggests even as late as 1859, years after the British had abolished the Transatlantic Trade, clandestine traders transported about 1000 Congolese captives into Puerto Rico (Chinea, 2016). In addition to forced migration from western Africa, prior to 1800, Puerto Rico’s proximity to the French and British sugar colonies in the Greater Antilles made it a site of frequent refuge for enslaved Africans escaping neighboring islands (Chinea, 1997).

Iberians who would have faced persecution in Spain, including North African Muslims, Sephardic Jews, and Christian Spaniards opposed to Spanish colonial control, were also drawn to Puerto Rico during the colonial period (Chinea, 1997). As an illustration of the cosmopolitan nature of Puerto Rico’s early colonial history, according to historical records, during the period between 1777 and 1797, over half of the island’s population was classified as non-white, with approximately 12% identified as enslaved Africans. As Puerto Rico’s importance as an agricultural and financially lucrative colony expanded in the late eighteenth century, this large non-white sector of the free population caused discomfort for white colonists and sparked widespread anti-Black sentiments, promoting ideas of *blanqueamiento* (literally, bleaching or whitening) that would persist into the centuries to come (Chinea, 1997; Picó, 2008). The effect of blanqueamiento is reflected across census taken from the late 19th century into the 21st century. The changing demographic indicates that the non-white population is reported to have declined throughout the latter half of the 19th century, with the most drastic decrease seen in the early 20th century with the non-white population consisting of 27% of the population (Vargas-Ramos, 2005).

Up until emancipation, enslaved peoples resisted and revolted against bondage in a variety of ways including escape, maroonage, and outright rebellion (Baralt, 2007). Post emancipation, Afro-Puerto Ricans continued to be vital to the Puerto Rican economy, laboring in a variety of sectors (Figueroa, 2005; Rodríguez-Silva, 2012). With the freedom of movement that came with emancipation, African descendants often moved on from locales where they had lived in bondage seeking new lives and opportunities in metropolitan centers, primarily in the south around Ponce and in the northeast in Loíza (Figueroa, 2005; Mayo Santana & Negrón Portillo, 2007). In 1898 Puerto Rico was formally ceded to the United States at the end of the Spanish American war and residents gained US citizenship in 1917 (Ayala, 2007). Many Afro-Puerto Ricans, like other Puerto Ricans, began to emigrate to US cities beginning in the late 19^th^ century (Laó-Montes, 2018). Periods of especially high migration from Puerto Rico to the US include the post-World War II era (1940-1960s), the financial crisis in the 2000s (2006-2016), and more recently, in the aftermath of Hurricane Maria (2017-present) (Hinojosa, 2018).

A variety of archivists have examined records and other documentation, including parish registers and government records, to glean information about the origins and experiences of African descendants in Puerto Rico. While some of these records provide insights, there are limitations to what may be interpreted from these historic sources. For example, enslaved Africans were often ascribed ethnicity corresponding to where they embarked for the Americas rather than how they self-identified. In addition, identifying information was not consistently collected from all disembarking peoples and historical records are temporally fragmented, providing perspectives on only limited periods of time (Stark, 2009). Moreover, beyond the perception that such alliances were threatening to Spanish settler-colonists, little is known about African and Indigenous relations, particularly within *palenques* (maroon communities) and other sites of resistance within Puerto Rico (Moscoso, 1995; Nistal-Moret, 2004).

The experiences of self-identified African descendants in Puerto Rico are further obscured by a history of anti-Blackness and racial discrimination. Puerto Rico, like several other former Spanish or Portuguese colonies in the Americas, has marketed itself for decades as a racial democracy, in which all inhabitants are assumed to be equally racially mixed. Everyone, in effect, is conceptualized as equally African, European, and Indigenous in ancestry (Godreau & Bonilla, 2021; Lloréns, García-Quijano, & Godreau, 2017). This idea of a universal *mestizaje* (mixture) was put in place in 1955 by the Institute of Puerto Rican Culture (ICP). The ICP was formed, in part, with the intention that Puerto Rico would develop a distinct national identity separate from that of the United States, one that would cast racism and racialization as ideas belonging uniquely to the U.S. This paradigm also explained away policies of *blanqueamiento* as simply a natural “whitening over time” that resulted from a society of racial harmony and mixture (Godreau & Bonilla, 2021). Furthermore, between 1950 and 2000 the question of racial identity was removed from the Puerto Rican census (Demby & Meraji, 2020). This notion that all Puerto Ricans are equally racially mixed, and that racial mixture has come to be equated with whiteness, thus renders Blackness on the island partially invisible and has produced a form of Puerto Rican nationalism that excludes Blackness. This leaves Afro-Puerto Ricans in a position where they cannot simultaneously legitimize their Blackness and their status as Puerto Ricans (Lloréns et al., 2017; Rivera, 2006). Individuals who self-identify as Afro-Puerto Rican therefore do so against a cultural landscape that has rejected the idea of their very existence.

Building on community engagement and support, we initiated this study with a specific goal of illuminating the experiences of self-identified Afro-Puerto Ricans, an understudied and minoritized population (Godreau, 2015). To address gaps in knowledge about Afro-Puerto Rican demographic history, we consider the genetic legacies of both African and European peoples within contemporary Afro-Puerto Rican communities. Genomic data, queried for genetic ancestry and admixture, provides a fuller assessment of the biogeographic origins of Afro-Puerto Ricans, particularly as they compare to additional populations in Puerto Rico and the wider Caribbean.

## 2. Materials and Methods

### 2.1 Ethics Statement

This project was a community engaged endeavor from the outset. Briefly, the project itself emanated from the community as many people were aware of previous genetic ancestry research done in Puerto Rico in the early 2000’s (Martinez-Cruzado et al. 2005). Despite the many insights gained about population genetics in Puerto Rico and other Caribbean communities, this body of work did not explicitly focus on historically Black-identifying communities. Consequently, many community members were interested in employing genomic technologies and welcomed this project as a means to address their questions about their own community history. Chapter 6 of Benn Torres & Torres Colón (2021, pp. 104–111) detail this initial community engagement. Additionally, local community members were incorporated into the project as hired consultants and fieldwork assistants for project recruitment.

Prior to engaging participants, all study protocols and materials were reviewed and approved by the Institutional Review Board (IRB) at Vanderbilt University (IRB protocol #170749). In addition, local collaborators and study coordinators reviewed and approved study goals and project design. All study participants provided written informed consent to participate in this project.

### 2.2 Sampling

Puerto Rican-born adults aged 18 or over, who self-identified as Afro-descendant or an equivalent term, and had at least three grandparents born in Puerto Rico were eligible to participate in the current study. A total of 58 individuals enrolled, providing saliva samples and completed genealogical interviews (Winful et al., 2023). During the interviews, participants were asked to list their birthplace, the birthplaces of their parents and grandparents, family surnames, and languages spoken within the home. Samples were collected at four sites on the island: in San Juan, Piñones/Loíza, Ponce, and Mayagüez (*Figure 1*). However, participants reported 13 different hometowns, stretching across the island. Reported hometowns include areas historically known or indicated by participants to be home to Afro-Puerto Rican communities.

**Figure 1:**
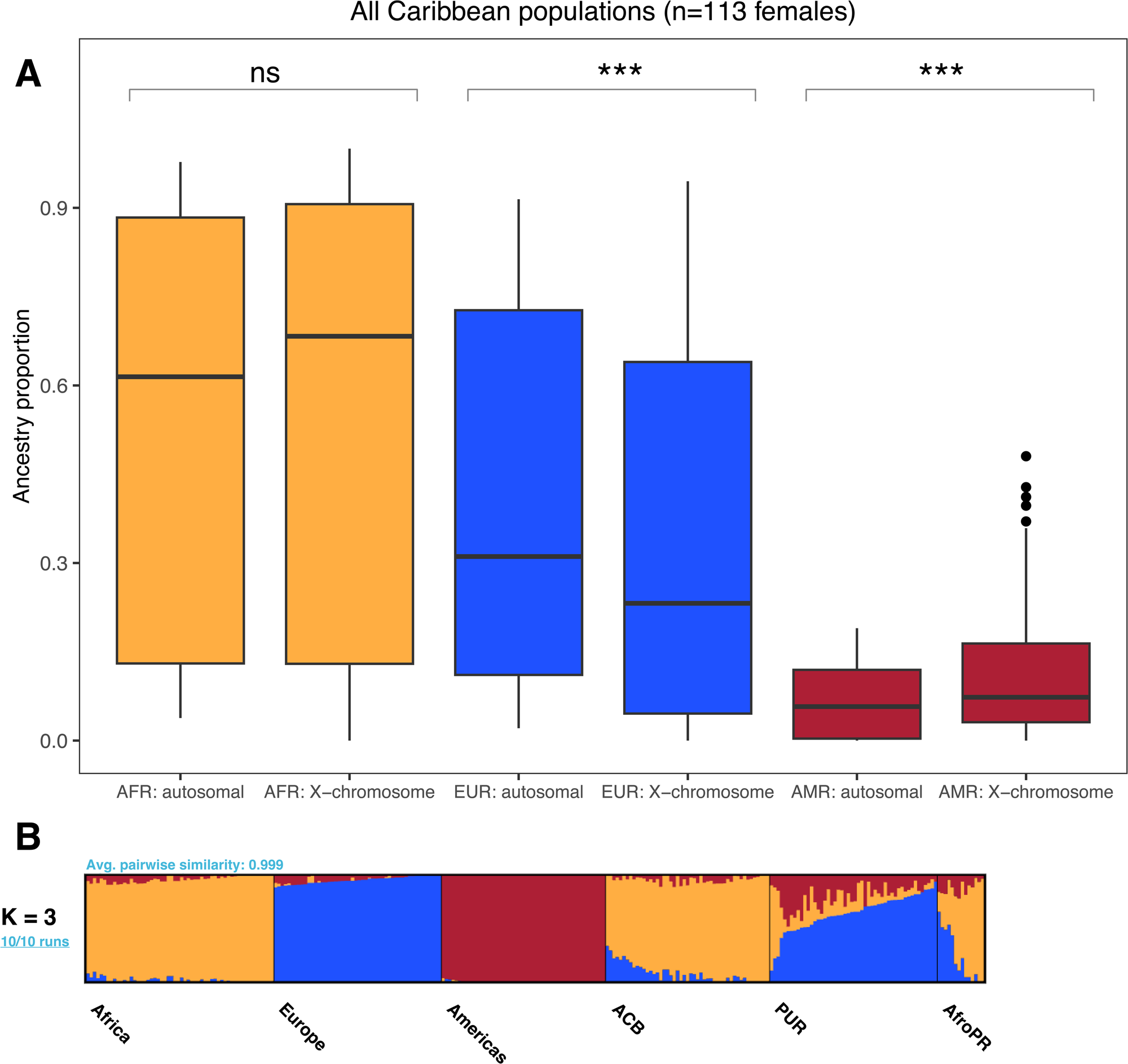
Map of study population and surrounding region. Circles denote approximate locations of sites where DNA samples were collected.

### 2.3 Sample Analysis

Upon providing consent and completing the interview, each participant provided 2 mL of saliva in an Isohelix Genefix™ (GFX-02) collection tube. Genomic DNA was then extracted using the Isohelix GeneFiX™ Saliva-Prep DNA Isolation Kit (GSPN) following manufacturer’s instructions. Each extracted sample was then checked for purity and quantified using a Nanodrop 1000 and Qubit® 3.0 Fluorometer, respectively. We recovered a median of 83 ng/μL genomic DNA from each sample. A subset (n=30) of the full sample was selected for autosomal analysis using National Geographic’s Genochip 2.0. These 30 samples were chosen based on the historical depth (amount of time the participant’s family had been located in Puerto Rico) and content of their associated genealogical interviews with the goal of elucidating more information about the African descendant experience in Puerto Rico.

The Genochip is a validated microarray containing thousands of genome-wide single nucleotide polymorphisms (SNPs) developed for the purpose of addressing questions about human origins and the global dispersion of *H. sapiens*. Various iterations of the Genochip were used during the Genographic Project (April 13, 2005-May 31, 2019), with the last microarray (Genochip 2.0) containing 750,000 genome-wide SNPs. Based upon Illumina’s OmniExpress microarray, the Genochip 2.0 was specially customized to maximize the number of genetic ancestry informative markers for global populations (Elhaik et al., 2013). The samples from the selected 30 individuals were genotyped at Family Tree DNA in Houston, Texas.

### 2.4 Quality Control

We used Plink 1.9 (Chang et al., 2015) to apply quality control measures to the genotype data prior to downstream analyses as recommended by Anderson et al. (2010). After ensuring that each sample met a 98% genotyping threshold we iteratively tested and, if necessary, removed SNPs or individual samples that failed to meet genotyping standards (Figure S1). Criteria for removing individuals or markers included sex assignment inconsistencies, missing data (>10% missing data for individual; >5% missing data for SNP), failed calls, or a failure to map to the GRCh37/hg19 human genome reference. After quality control, our final dataset contained 662,669 SNPs for each of the 30 participants. This final dataset included only autosomal markers (chromosomes 1-22), excluding all mitochondrial and sex chromosome SNPs. An additional dataset including 16,764 X-chromosome SNPs from all 30 participants was also prepared from the quality controlled files for further analysis.

### 2.5 Data Analysis

Using Plink 1.9, we merged the Afro-Puerto Rican dataset with reference data from four populations from the 1000 Genomes Project (1KGP) Phase 3 (2015): Yoruba (YRI, n=101), Northern European (CEU, n=93), Puerto Ricans (PUR, n=104) and Afro-Caribbean from Barbados (ACB, n=96) (Sudmant et al., 2015; The 1000 Genomes Consortium, 2015). Lastly, we included genotype data from 100 Peruvians from Puno with large proportions of Indigenous American ancestry previously published as part of the Population Architecture Using Genomics and Epidemiology (PAGE) study (Wojcik et al., 2019). For consistency, all reference datasets were QC filtered as detailed above (Figure S1).

To examine the relationship between sampled Afro-Puerto Ricans and putative parental (reference) populations, we first implemented SmartPCA from the EIGENSOFT package (Patterson, Price, & Reich, 2006; Price et al., 2006). Prior to running the analysis, the merged dataset was LD filtered (r^2^ threshold of 0.1 for over a window size of 50 and step size of 10), resulting in a subset dataset of 50,801 SNPs. The SmartPCA results were then plotted using R version 4.2.2 and the R ggplot2 package (R Core Team, 2020; Wickham, 2016). Clustering analyses were then completed in ADMIXTURE (Alexander & Lange, 2011) using the LD-pruned merged dataset. We performed 10 replicates of ADMIXTURE analysis for models associated with K=2 through K=10 clusters to assess the relative likelihood of any given K-way ancestral model. The aggregate ADMIXTURE results were plotted using Pong (Behr, Liu, Liu-Fang, Nakka, & Ramachandran, 2016).

We additionally analyzed data from chromosome X in isolation in order to investigate the presence of sex-biased admixture among the Afro-Puerto Rican and 1KGP ACB and PUR populations. As diploid chromosomes are required for this analysis, only chromosomal females were included (AfroPR: n=14, PUR: n=50, ACB: n=49, YRI: n=56, CEU: n=50, Puno: n=49). After LD-pruning, the dataset had 1271 SNPs. We followed the same procedure for running ADMIXTURE on the X chromosome data as described above for the autosomal data. We then compared X chromosome and autosomal ancestry proportions for the same individuals with the Wilcoxon signed-ranks test as in Moreno-Estrada et al. (2013). This test is a non-parametric alternative to a t-test that assesses whether significant differences exist between two distributions (Moore, McCabe, & Craig, 2007). The null hypothesis is that the distribution of ancestry proportions on the autosomal chromosomes is identical to the X chromosome. The test was conducted using the R pairwise.wilcox.test function at several significance cutoffs (p<0.05, p<0.01, p<0.001) and using the Bonferroni multiple tests correction. The distribution of ancestry proportions across populations was visualized with boxplots.

To perform subcontinental analyses for the European and African ancestral components of the Afro-Puerto Rican individuals, we implemented the ancestry-specific multidimensional scaling (MDS) approach published by Browning et al. (2016). This method performs an MDS on a Euclidean distance matrix, which therefore produces principal components as in a PCA (Cox & Cox, 1994; Gower, 1966). For these analyses we constructed two subcontinental reference panels of previously published genotype data. The African subcontinental panel included 44 African populations merged from the 1KGP Phase 3, African Genome Variation Project (Gurdasani et al., 2015) and Patin et al. (2017). The European subcontinental panel included 19 European and diasporic Jewish populations merged from the 1KGP Phase 3, Human Genome Diversity Panel (HGDP) (Bergström et al., 2020), Behar et al. (2010), and Hernández et al. (2020) datasets (Table S3). For each panel, we followed the population classification scheme presented in the original publications. We filtered these datasets according to the same QC parameters used to prepare the sample data. We next merged these subcontinental reference panels with the Afro-Puerto Rican dataset, the 1KGP PUR (n=104) and ACB (n=96) populations, and three continental reference populations from the 1KGP (CEU, n=93; YRI, n=101; and PEL, n=83) using Plink 1.9. Although we found the PAGE Puno cohort to be a suitable Indigenous American reference population for the ADMIXTURE and SmartPCA analyses, we used the 1KGP Peruvians from Lima (PEL) cohort for the RFMix and MDS analyses that follow due to higher SNP overlap with our study and reference populations allowing for more robust analysis of subcontinental ancestry (Figure S2). In total, the subcontinental African combined panel includes 164,851 SNPs and 2,524 individuals. The subcontinental European combined panel includes 205,312 SNPs and 1,139 individuals.

We then prepared these panels for MDS analysis using the steps and scripts provided by Browning et al. (2016) (https://faculty.washington.edu/sguy/local_ancestry_pipeline/). Following the structure of Browning et al.’s analysis, we used Plink 1.9 to prepare per-chromosome VCF files from the final merged datasets. We then used Beagle 4.0 (Browning & Browning, 2007) to phase these VCF files and prepare them for local ancestry estimation with RFMix 1.5.4 (Maples, Gravel, Kenny, & Bustamante, 2013). At this point, we also prepared RFMix input files used to indicate which haplotypes are associated with admixed populations (and therefore need local ancestry estimation), as well as the allelic variant data and SNP locations. RFMix and the following MDS analyses are performed on a per-haplotype rather than per-individual basis (N=2*individuals) due to possible variation in ancestry between haplotypes within each individual. To verify the accuracy of the RFMix ancestry estimation, we followed Browning et al (2016) and recorded the highest Forward Backward posterior probabilities for each ancestry per site at each haplotype. We report the autosomal averages for both the African and European combined panels in Table S4. Regardless of the ancestry assigned, the Forward Backward posterior probability is close to 1 (between 0.97 and 0.99) for the combined European subcontinental panel. For the combined African subcontinental panel average posterior probabilities are also close to 1 for the African (0.985) and European ancestry calls (0.955), but 0.69 for the Indigenous American ancestry calls. Lower accuracy is likely due to the lower SNP density of this panel and lower proportions of Indigenous American ancestry represented in the studied populations. Overall, average local ancestry estimation posterior probabilities are 0.987 and 0.879 for the European and African subcontinental panels, respectively.

Once RFMix was run for the admixed populations, we used Beagle 4.0 to combine all chromosomal output into a single file and filtered individual haplotypes based on their RFMix-estimated ancestry proportions. Following the direction of Browning et al., we LD filtered the dataset after performing RFMix, to avoid the creation of spurious PCs due to the inclusion of genomic regions in high linkage disequilibrium. Similar to the SmartPCA and ADMIXTURE analyses, we used an r^2^ threshold of 0.1 for pruning over a window size of 50 and a step size of 10. After LD filtering, we were left with 82,388 SNPs for the analysis concerning the African ancestry component, and 62,681 SNPs for the analysis of the European ancestry component (Figure S2).

MDS analyses were performed using the dist and cmdscale functions in R, utilizing the script provided by Browning et al. We performed the MDS analysis for four ancestry thresholds, removing haplotypes that did not have at least 10%, 15%, 20%, and 25% of European or African ancestry, respectively in the admixed populations (Table S5). These data preparation steps and subsequent analyses were performed on the Minnesota Supercomputing Institute’s (MSI) agate cluster. Results were plotted using R and ggplot2. Note that because the MDS method utilized here returns a fixed number of principal component axes, we are unable to calculate the amount of variation explained by each axis. The reference populations in the European and African subcontinental panels were organized by geographic regions and ethnic group identification, in order to best highlight relevant historical classification of populations and highlight genetically distinct or isolated populations (Table S3)

Scripts used for data quality control, preparation, and analysis are located at the public github repository https://github.com/eculrich/AfroPR_Scripts.

## 3. Results

### 3.1 Continental Affinities in Afro-Puerto Ricans

Our initial analysis using SmartPCA addressed the question regarding continental genetic ancestry within Afro-Puerto Ricans. The first three components of the PCA plot explained about 11% of the variation within the data (*Figure 2*). The first component, PC1, separated African from non-African populations, PC2 separated Indigenous American populations from all other populations, and PC3 separated European populations. When plotting PC1 versus PC2, Afro-Puerto Rican individuals clustered between the Puerto Rican (PUR) and Afro-Caribbean (ACB) reference populations in a clinal pattern which itself was nested between European (CEU) and African (YRI) reference populations. In the PC1 versus PC3 plot (Figure S3b), individuals clustered based on broad geographic regions. All the Puerto Rican individuals (both reference and the study populations) tend to be distributed clinally between the African and European reference groups. Within this clinal pattern, the reference Puerto Rican individuals are generally near the European reference group while the Afro-Puerto Ricans are generally centrally located, widely dispersed between African and European reference groups. The ACB reference group clusters most closely to the African reference population. PC2 versus PC3 (Figure S3a, Figure 2a), has a similar distribution as PC1 versus PC3, with Afro-Puerto Ricans clustering between the African and European reference groups. In each PC plot, Afro-Puerto Ricans exhibited a fair amount of heterogeneity, with some Afro-Puerto Rican individuals showing great affinity with African and African-descendant reference populations, while other Afro-Puerto Ricans appeared more similar to reference 1KGP Puerto Ricans.

**Figure 2:**
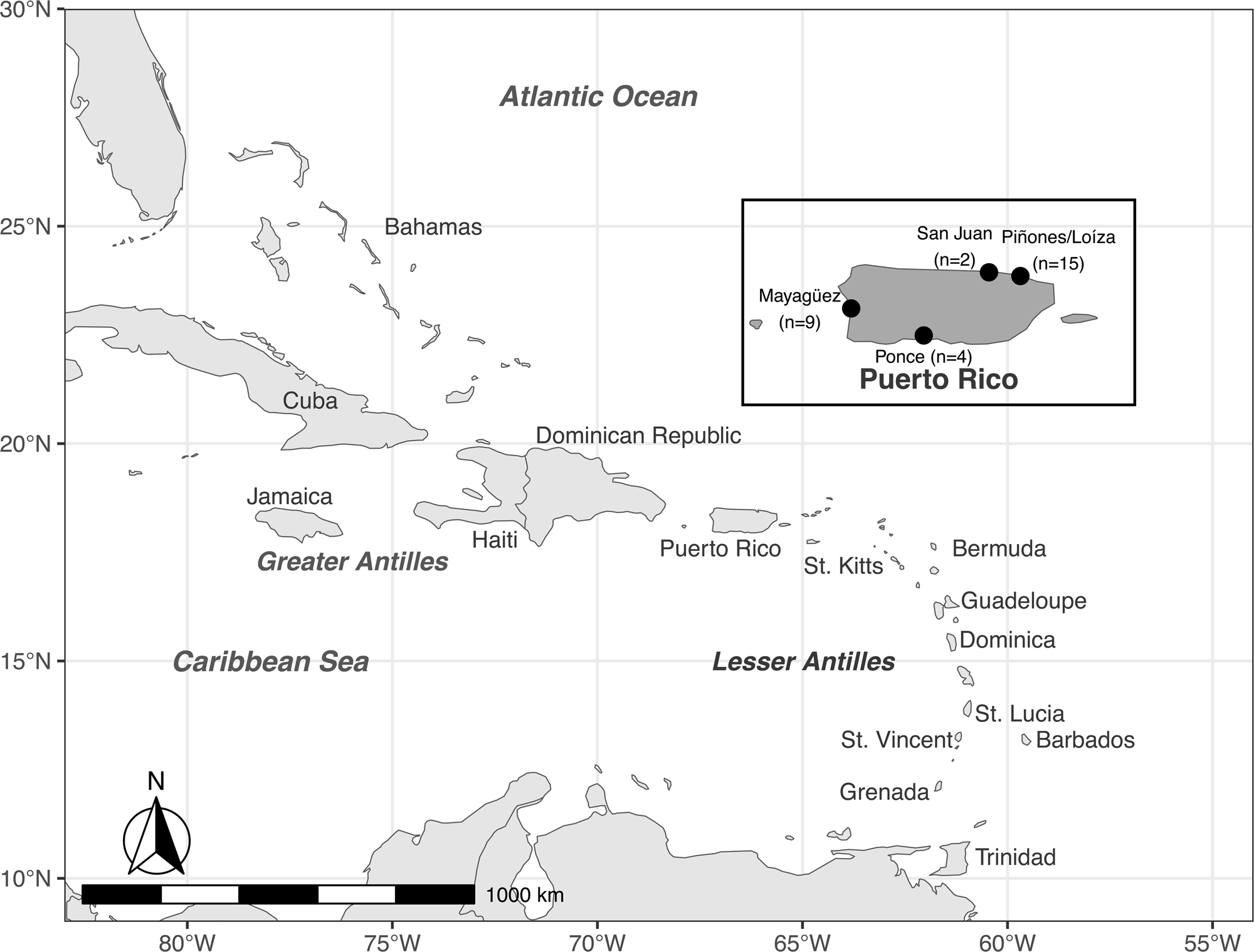
Population structure of study population and comparative regional populations. (A) Principal components analysis of AfroPR sample individuals (represented by open circles), two 1000 Genomes Project Caribbean populations (PUR and ACB, represented by open circles and crosses, respectively), and three 1000 Genomes Project continental references (YRI, CEU, PEL, represented by colored circles). (B) ADMIXTURE analysis for the same populations featured in A showing an unsupervised clustering model assuming K=3. Thin vertical bars represent individuals, with black bars separating populations. At K=3, the clusters correspond roughly to continental reference populations (African, European, and American), with Caribbean populations demonstrating a mix of continental affinities.

### 3.2 Admixture in Afro-Puerto Rican Communities

We estimated global genetic ancestry proportions to consider the impacts of continental populations on the genetic variation of Afro-Puerto Ricans. According to the ADMIXTURE analysis inclusive of the reference populations, three clusters returned the smallest error (CV error=0.54355) (*Figure S4*). Thus, a three-way admixture model best represents global continental contributions to Afro-Puerto Rican populations (*Figure 2b*). This result is consistent with the findings of previous research based on mtDNA (Martínez-Cruzado et al., 2005; Vilar et al., 2014; Winful et al., 2023) and autosomal (Bryc et al., 2010; Moreno-Estrada et al., 2013) datasets. The Afro-Puerto Rican individuals primarily displayed genetic ancestries similar to reference populations from Africa (average genetic ancestry proportions = 58.8%, SD=17.99%), Europe (34.5%, SD=16.04%), and the Americas (6.7%, SD=2.4%). Relative to the other Caribbean reference groups, Afro-Puerto Ricans had less African continental genetic ancestry than ACB (87.8%, SD=7.73%) and more African continental genetic ancestry than PUR (14.8%, SD=9.8%). European continental genetic ancestry was highest in the PUR reference group (73.11%, SD=10.27%) and lowest in the ACB reference population (11.6%, SD=7.4%), while genetic ancestry from the Americas was the highest in the PUR reference group (12.1%, SD=3.54%) and lowest in the ACB population (0.55%, SD=1.02%) relative to the Afro-Puerto Rican sample (Table S1).

To identify instances of sex-biased admixture among Afro-Puerto Ricans, genetic ancestry proportions were characterized on the X chromosome separately from the autosomes. To ensure that only diploid loci were compared, this analysis was performed only on chromosomal females (see Materials and Methods). As with the autosomes, ADMIXTURE analysis on the X chromosome dataset found K=3 to be the best model fit (CV error = 0.53385) *(Figure S5).* Comparisons of autosomal versus X chromosome genetic ancestry proportions revealed no significant differences in proportions of African ancestry between loci (p>0.05), but found a significant increase in Indigenous American genetic ancestry, and a reduction in European genetic ancestry on the X chromosome for all three Caribbean populations (p<0.05) *(Figure 3, Table S2)*.

**Figure 3:**
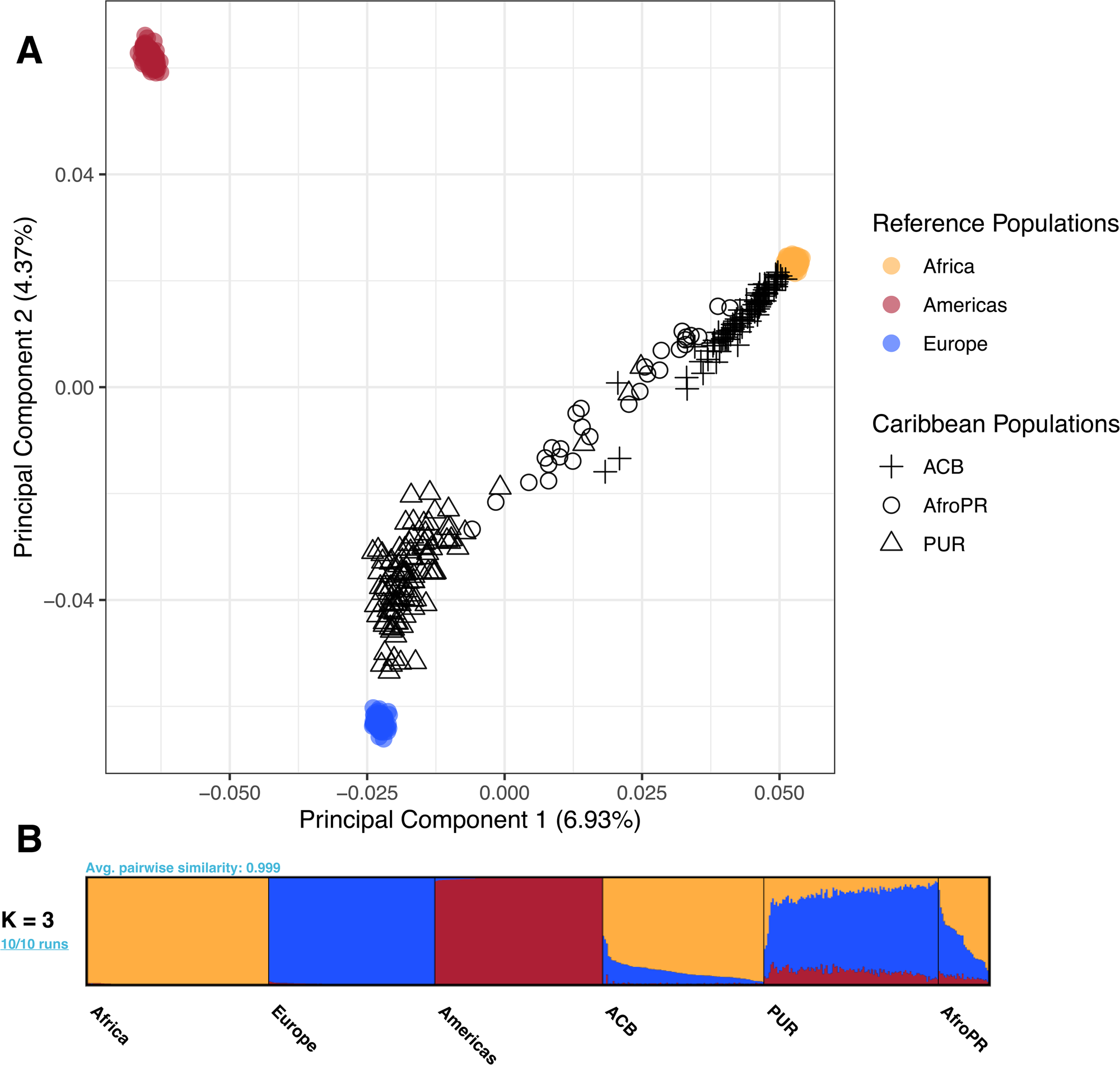
Comparison of population structure on the X versus autosomal chromosomes for the study population and comparative regional populations. (A) Boxplots with associated Wilcoxon-rank sum calculated significance values comparing African, European, and American ancestry proportions on the X chromosome and all autosomal chromosomes for chromosomal females from the study population and two 1000 Genomes Project Caribbean populations (PUR, ACB). Ancestry proportions were estimated using ADMIXTURE. No significant difference was found for the African ancestry proportion between X chromosome and autosomal data, but a significance corresponding to p ≤ 0.001 is associated with the European and American ancestry proportions. (B) ADMIXTURE analysis for female X chromosomes from five 1000 Genomes Project continental reference and Caribbean populations and the study population, assuming a K=3 model. Thin vertical bars represent individuals, with black bars separating populations. At K=3, the clusters correspond roughly to continental reference populations (African, European, and American), with Caribbean populations demonstrating a mix of continental affinities.

### 3.3 West African Genetic Ancestry in Afro-Puerto Ricans

Beyond considering global genetic ancestry, we also focused specifically on the African ancestral component to elucidate which subcontinental regions in Africa contributed genetic ancestry to contemporary Afro-Puerto Ricans. In the ancestry-specific **MDS** plot with a 20% African genetic ancestry threshold (*Figure 4a)*, the first two components reflected the geographic pattern of West and Central Africa, moving from the Senegambian region in the north, southward to the Bight of Biafra, then eastward toward Central Africa and Angola.

**Figure 4:**
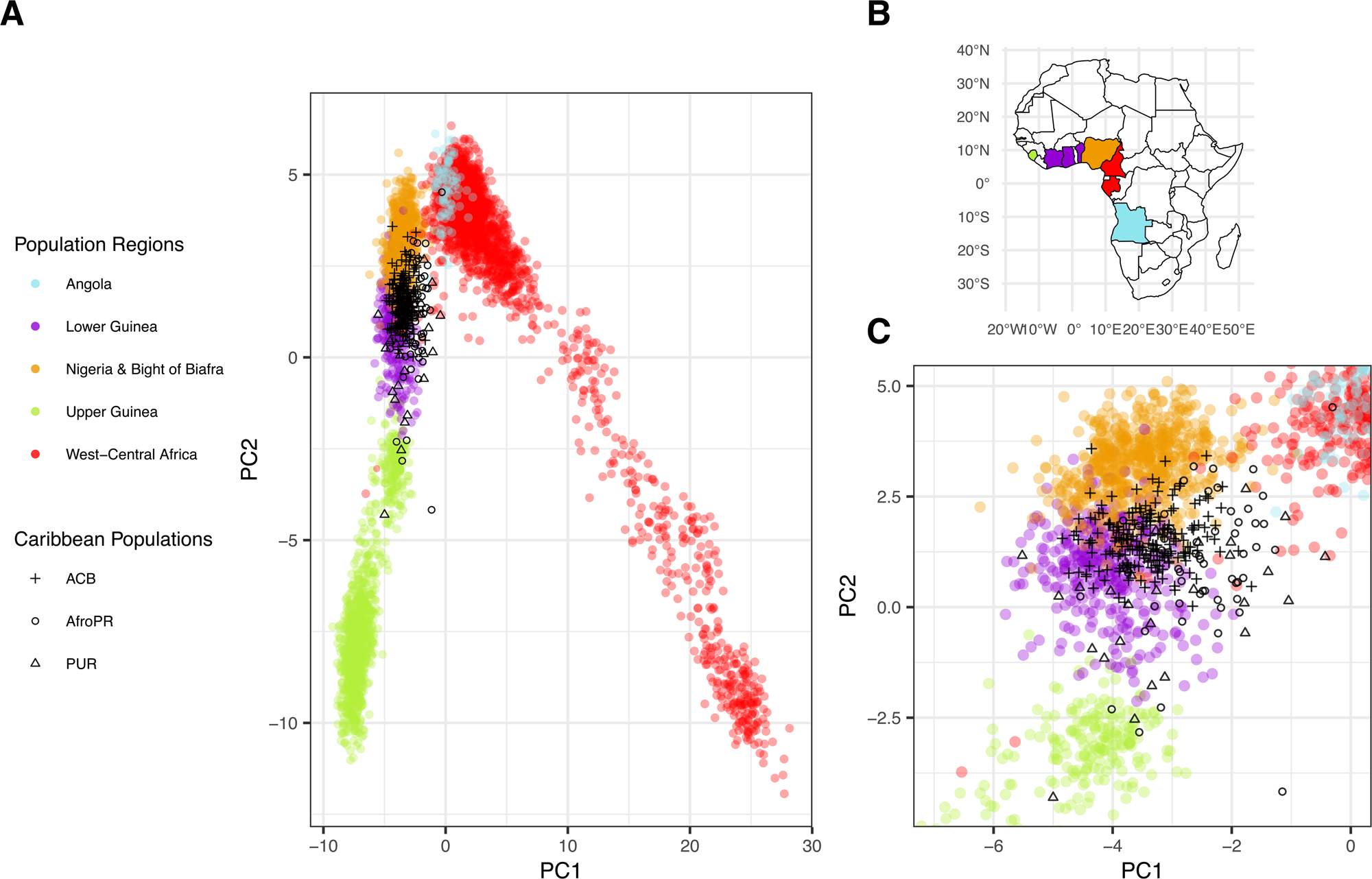
Ancestry-specific MDS analysis of the African ancestry of the study population and comparative regional populations. (A) MDS analysis of individual haplotypes with at least 20% African ancestry calculated by RFMix. Forty four populations from coastal regions of Africa corresponding to possible historical sources of African ancestry in the Caribbean (colored circles) are plotted alongside the study population (open circles) and two 1000 Genomes Project Caribbean populations (ACB: crosses, PUR: open triangles). At a 20% ancestry cutoff, 58/60 study population haplotypes, 192/192 ACB haplotypes, and 22/208 PUR haplotypes are included. (B) Map of Africa including countries of origin for the selected African reference populations. Colors on the map correspond to the regional labels used in Panel A. (C) Zoomed-in region of the plot shown in Panel A featuring most of the Caribbean haplotypes. At this level of magnification, affinities are most apparent between the Caribbean populations and selected Nigerian and Lower Guinean reference populations.

Within this plot, the Afro-Puerto Ricans were dispersed from the northern regions to the southeastern regions of West Africa, showing the most affinity with reference populations from Lower Guinea, Nigeria and the Bight of Biafra, and to a lesser extent Upper Guinea and West Central Africa. Unsurprisingly, the PUR reference group also showed the same African ancestral components as were observed in Afro-Puerto Ricans. The distribution of Afro-Puerto Rican individuals in the plot was somewhat different from the distribution of reference ACB individuals who were primarily concentrated with reference populations from Lower Guinea, Nigeria and the Bight of Biafra, illustrating a more homogeneous African ancestral component relative to Afro-Puerto Ricans. These broad patterns are maintained irrespective of the genetic ancestry thresholds used (10%, 15%, 20%, 25%), but at higher thresholds the number of PUR individuals removed from the analysis also increases due to their overall lower levels of African genetic ancestry (*Figure S7).* Overall, this plot illustrates that Puerto Ricans in general show a wider distribution of African ancestral affinities than that observed in Afro-Caribbeans from Barbados.

### 3.4 European Genetic Ancestry in Afro-Puerto Ricans

We next focused specifically on the European genetic ancestral component of contemporary Afro-Puerto Ricans. In the ancestry-specific **MDS** plot with a 20% European genetic ancestry threshold (*Figure 5)*, the first two components reflect the geographic pattern of Europe, moving from Central and Eastern Europe to the Iberian peninsula. Populations with unique genetic substructure, such as the Basque, Sardinians and diasporic Jews, cluster separately from other Europeans (Behar et al., 2010; Chiang et al., 2018; Flores-Bello et al., 2021; Marcus et al., 2020; Ostrer & Skorecki, 2013). Within this plot, Afro-Puerto Ricans and the PUR reference group clustered primarily with Iberian and Italian populations, although some individuals showed more affinity with reference populations from the Basque country and with Sardinians. The distribution of Afro-Puerto Rican and PUR individuals in the plot was different from the distribution of ACB individuals who were primarily concentrated with reference populations from Western Europe. This pattern is replicated irrespective of genetic ancestry threshold, but at higher thresholds the number of ACB and Afro-Puerto Rican individuals removed from the analysis increases due to their lower levels of European genetic ancestry *(Figure S10).* Overall, this plot illustrates that European ancestral affinities differ between Puerto Ricans in general and Afro-Caribbeans from Barbados.

**Figure 5:**
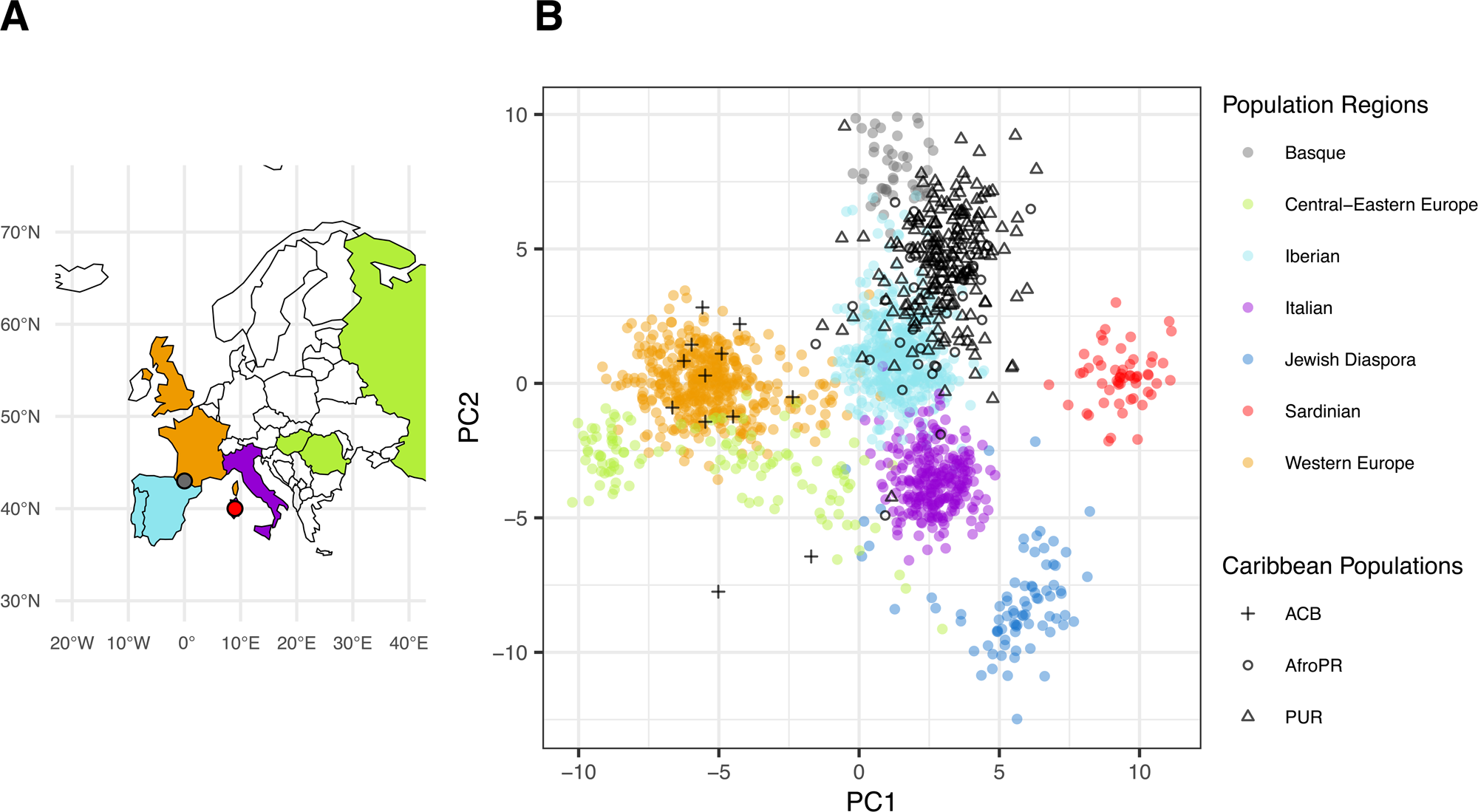
Ancestry-specific MDS analysis of the European ancestry of the study population and comparative regional populations. (A) MDS analysis of individual haplotypes with at least 20% European ancestry calculated by RFMix. Nineteen populations from Europe corresponding to possible historical sources of European ancestry in the Caribbean (colored circles) are plotted alongside the study population (open circles) and two 1000 Genomes Project Caribbean populations (ACB: crosses, PUR: open triangles). At a 20% ancestry cutoff, 40/60 AfroPR haplotypes, 12/192 ACB haplotypes, and 208/208 PUR haplotypes are included. (B) Map of Europe including countries and regions of origin for the selected European reference populations. Colors on the map correspond to the regional labels used in Panel B.

Finally, we note that given the low proportions of Indigenous American autosomal ancestries identified in the three admixed Caribbean populations in this study (average genetic ancestry proportions = Afro-Puerto Ricans: 6.7%, PUR: 12.1%, ACB: 0.55%), and lower accuracy of local ancestry estimation for Indigenous American chromosomal segments, we were unable to run ancestry-specific MDS analyses for the Indigenous American ancestral component with robust confidence. We decided upon a minimum threshold of about 15% for genetic ancestry specific MDS based on the precedence set in similar work (Rodríguez-Rodríguez et al., 2022).

## 4. Discussion

In the current study we sought to illuminate the demographic histories of self-identified Afro-Puerto Ricans, with specific consideration to the genetic legacies of African and European ancestors in shaping the contemporary population. The principal component analyses showed that Afro-Puerto Ricans have genetic ancestry from African, Indigenous American, and European peoples. The global genetic ancestry estimates quantify these findings, illustrating that most Afro-Puerto Ricans have primarily African genetic ancestry, followed by European genetic ancestry, and smaller but substantial amounts of Indigenous American genetic ancestry. Furthermore, the PCA plots (Figure 2, S3) indicate that genetic ancestry from these continental populations is not homogeneous, rather, given the distribution of Afro-Puerto Rican individuals within the plot, amounts of continental genetic ancestry vary substantially between Afro-Puerto Rican individuals. These results are congruent with previous genetic and genomic studies of the general Puerto Rican population (Bertoni, Budowle, Sans, Barton, & Chakraborty, 2003; Fernandez Cobo et al., 2001; Via et al., 2011; Vilar et al., 2014). While these published studies do not specifically focus on self-identified Afro-Puerto Ricans, they all show that genetic ancestry among Puerto Ricans can be traced to these three continental populations. Accordingly, these genomic findings also corroborate with archival, archeological, and historical accounts of the origins of contemporary Puerto Rican society (Ayala, 2007; Curet, 2005; Picó, 2008).

Our analyses also revealed a significant skew towards a higher proportion of Indigenous American genetic ancestry, and a lower proportion of European genetic ancestry in the X versus autosomal loci in all three Caribbean populations (AfroPR, PUR and ACB). These results point to the presence of strong sex bias during the colonization of the Caribbean islands, a pattern previously shown by earlier genetic studies using both uniparental and autosomal markers (Adhikari, Chacón-Duque, Mendoza-Revilla, Fuentes-Guajardo, & Ruiz-Linares, 2017; Moreno-Estrada et al., 2013; Nieves-Colón, 2022). Historical documents demonstrate that although Iberian women did participate in the early phases of Spanish colonization, they initially came to the Caribbean in much lower numbers than male settlers. As the formation of families was encouraged by the Spanish Catholic Church, male settlers took (sometimes forcibly) Indigenous women or *ladinoized* (Christianized) African women as wives, concubines or domestic partners (Altman, 2013). However, while Indigenous-European *mestizo* children were quickly absorbed into Puerto Rican colonial society, children that had one African and one European parent were viewed as peripheral, highly associated with enslavement and low status, a view that has influenced the perception of Afro-Puerto Rican identity into the present (Schwartz, 1997).

Regarding the African genetic ancestral component, ancestry-specific MDS analysis indicated that Puerto Ricans have African genetic ancestry from a wide swath of western Africa, specifically, Upper and Lower Guinea and west Central Africa. This finding most likely reflects the different iterations of African influx into the island. Archival records detailing the geographic origins of enslaved Africans recorded the aforementioned regions as sources of enslaved labor imported into Puerto Rico. These types of records however, cannot be accepted entirely on face-value, since the ethnic and geographic origins of enslaved persons was not consistently nor systematically collected by enslavers (Hall, 2005). According to a study of baptismal documents, historian David Stark (2009) notes that parish records dating to the 16^th^ century indicate that Guinea Bissau, the Bijagos islands, Senegal (classified as Upper Guinea in our AS-MDS analysis) and also Ghana were primary sources of enslaved labor due to their navigationally advantageous geographical positions and the expertise in cattle ranching of enslaved peoples from these areas. Moving into the 17^th^ century, enslaved peoples tended to come from Congo and Angola as a result of shifting economic and political alliances within the ruling Spanish class. In the 18^th^ century, slave trafficking into Puerto Rico by French and British traders shifted trade back to peoples from Upper Guinea and regions of west central Africa (Stark, 2009; Wheat, 2016, p. 29).

In addition, the geographic distribution of ancestries differed from what was detected for the other comparative Caribbean population from Barbados. Unlike the Puerto Rican individuals that were spread throughout the plot, ACB individuals tended to concentrate with populations from Lower Guinea and the Bight of Benin (Figure 4). While additional research is needed to identify why we see this particular pattern, it is suggestive that either the sampling of 1KGP ACB is limited and not representative of the island, or that Puerto Rico and Barbados had very different histories with regard to the nations and communities that were the involved in slave trade and inter-island movement. Furthermore, we can see that the Afro-Puerto Ricans have a wider distribution in this plot relative to the ACB individuals. These findings suggest there may be some unrecognized heterogeneity of African descendants throughout the Caribbean.

Previous studies of African genetic ancestry in the Caribbean have focused largely on Anglophone Afro-descendent populations or have not distinctly considered Afro-descendent populations in primarily Spanish-speaking islands. Despite this dissimilarity in population focus, our finding of heterogeneity in African genetic ancestry throughout the Caribbean has been previously documented in both mtDNA and autosomal DNA studies (Benn Torres, Kittles, & Stone, 2007; Bukhari, Luis, Alfonso-Sanchez, Garcia-Bertrand, & Herrera, 2017; Gouveia et al., 2020; Moreno-Estrada et al., 2013; Stefflova et al., 2011). While we did not include ancient DNA comparisons for historic populations, previous studies of ancient DNA of enslaved individuals in the Caribbean also suggest early heterogeneity with respect to African origins (Schroeder et al., 2015). Our finding that Afro-Caribbeans from Barbados have less varied sources of African genetic ancestry relative to the Puerto Rican populations has previously been identified by studies that suggest that Spanish-speaking colonies in the Americas tended to have more heterogeneity with respect to their African genetic ancestry, possibly because Spain did not have designated colonies in Africa (Gouveia et al., 2020; Stefflova et al., 2011). Additionally, previous studies have largely identified genetic affinity with Nigerian and Yoruba populations (Montinaro et al., 2015; Murray et al., 2010). These results have been questioned by Jackson (2021) as being influenced by an overrepresentation of Yoruba and Nigerian genomes in available African genetic datasets, which may be masking more complex patterns of African ancestry in the Americas.

Ancestry-specific MDS further indicates that both Afro-Puerto Ricans and the PUR population share genetic ancestries with reference populations from Iberia and southern Europe. These patterns are consistent with the historical record which describes at least two major pulses of European migration into Puerto Rico during the colonial period: the initial European invasion during the 15th and 16th centuries, and a second wave during the 19th century when the Spanish Crown promoted the migration of European Catholics into its American colonies.

Incentivized by the Royal Decree of Grace (1815), which promised settlers wealth, land and access to enslaved labor, the growing colonial economy of Puerto Rico attracted large numbers of migrants from the Balearic and Canary Islands, Corsica, and France (Chinea, 1996). Although in lesser numbers, settlers from other European powers such as Britain, France, Belgium and Denmark also migrated to Puerto Rico during this time (Cifre de Loubriel, 1964; Knight, 2003; Rodríguez Mendoza, 2004; Santana Pérez, 2008). Moreover, as wars of liberation exploded in the continental Americas, creole elites from neighboring colonies such as Haiti, Cuba and Venezuela also relocated to Puerto Rico (Cifre de Loubriel, 1964; Picó, 2008). Many of these settlers came with their families, establishing a new merchant and landowner class that clustered around sugar and coffee estates in central and southern Puerto Rico (Scarano, 1981). In contrast, ACB individuals clustered closely with reference populations from Western Europe, indicating differences in the subcontinental European origins of African descendants across the Caribbean.

Our findings are also consistent with previous genetic studies which have found that the dynamics of colonization structured the distribution of Eurasian ancestries found across the Caribbean and broader Latin America (Adhikari et al., 2017; Mendisco et al., 2019; Montinaro et al., 2015; Nieves-Colón, 2022). Several studies examining European genetic ancestry in the Caribbean have identified a high degree of affinity with contemporary Iberian populations, particularly among Spanish-colonized islands (Fortes-Lima et al., 2017; Montinaro et al., 2015; Moreno-Estrada et al., 2013). Other studies have identified genetic ancestry components in Puerto Ricans contributed by populations that we were unable to include as references here due to low levels of SNP overlap, including North Africans and Canary Islanders (Díaz-Zabala, Nieves-Colón, & Martínez-Cruzado, 2017; Shi et al., 2023), indicating potential comparisons for future studies. Additionally, Jewish *converso* genetic ancestry has been identified in Latin America (Chacón-Duque et al., 2018), and historical documents suggest that *conversos* were present across Ibero-America, including in Puerto Rico (Alicea Rivera, 2017; Kagan & Morgan, 2009). Interestingly, our study did not find a strong affinity between contemporary diasporic Jewish populations, including Sephardic Jews from Iberia, and Puerto Rican populations, but population labels such as “Iberian” or “Spanish” from studies that did not consider Jewish *converso* genetic ancestry may be overlooking the presence of contemporary Jewish or Jewish-descent populations in Iberia.

While the current study is consistent with previous work on Puerto Rican populations, the specific focus on a Puerto Rican subpopulation, self-identified Afro-Puerto Ricans, allows for the additional consideration of the intertwining of biology and culture regarding genetic ancestry and identity. As detailed in previous work (Benn Torres, 2014, 2018, 2020), the relationship between genetic ancestry and (social) identity is not direct, but rather a complex interplay of historical, political, and socioeconomic factors. How individuals choose to socially identify may or may not be influenced by knowledge of their genetic ancestry; what remains consistent however, is that social identifiers such as race are not biological (e.g. genetic) in nature but rather experiential social constructs. Furthermore, how biological information is interpreted and informs social identities varies across time and space (Benn Torres & Torres Colón, 2021).

In the current study, the genetic analyses provide insights into how genetic ancestry potentially functions within Puerto Rican communities. Both the PCAs and the ADMIXTURE analyses align with the idea of a Puerto Rican national narrative of *mestizaje*. In this narrative, Puerto Ricans trace their ancestries to African, Indigenous American, and European peoples, resulting in a variety of consequences in relation to political, economic, and social factors (Demby & Meraji, 2020; Godreau, 2015; Godreau & Bonilla, 2021; Rivera-Rideau, 2013; Rodríguez-Silva, 2012). Iin this cited work, authors challenge the ideas of racial fusion and harmony that have been perceived as fundamental to the Puerto Rican tripartite ancestry narrative. Moreover, Puerto Rico’s history in relation to the US, inclusive of ongoing migration to and from the US mainland, also bears some influence on ideas about Puerto Rican identity (Godreau & Bonilla, 2021).

As discussed in the Introduction, the population central to this study consists of individuals who have self-identified as Afro-Puerto Rican within this culture of universal *mestizaje* and an aura of discomfort surrounding notions of race. In this context, scholar Isar P. Godreau puts forth the idea of racial “scripts” within Puerto Rico that dictate how and where Puerto Rican Blackness can exist (2015). The predominant racial script within Puerto Rico is that because Puerto Rico did not start as a plantation colony, few slaves lived on the island, leading to a whitening over time of their mixed-race descendants. The minimal African influence, according to this narrative, can still be felt in aspects of culture such as music, spirit, and rhythm, found in all Puerto Ricans. Coastal communities that are largely Black have preserved these manifestations of Blackness despite modernization and Americanization, denying the role of *blanqueamiento* in the relative whitening of many other Puerto Rican communities. This narrative, much like the narrative surrounding some of the earliest Black and European mixed inhabitants of the island, equate Blackness with slavery and deny that people of color in Puerto Rico may be considered Black (Godreau, 2015; Schwartz, 1997). As such, “Blackness on the island is equated with slavery, inferiority, awkwardness, ignorance and being uneducated, comical, hypersexual, and of the underprivileged class”, denying avenues for Blackness to be praised or acknowledged outside of certain spatial or stereotypical facets (Rivera, 2006, p. 167). Because of the geographic and spatial aspect of this racial script or narrative (that Black communities are largely coastal and rural), there is an additional underlying script that individuals beyond this space cannot identify themselves as Black Puerto Ricans. The act of binding Blackness to a place, especially one that represents a stronghold against modernization or Americanization, denies individuals outside of these places the legitimacy of their Blackness and Puerto-Ricanness (Godreau, 2015).

In effect, these ideas of racial democracy and the permissible scripts of Blackness in Puerto Rico result in a state that defines itself as overwhelmingly white or Latino rather than Black. In the 2010 census, 74% of the Puerto Ricans surveyed identified themselves as only white rather than mixed race (Lloréns et al., 2017). In the more recent 2020 census, only 7% of the population identified as “Black or African American alone”. This value is a 50.4% decline relative to the proportion of people that responded that way in the 2010 census (Figueroa-Lazu, Hinojosa, & Bonilla, 2022) This decline may be potentially explained by increases in the proportion of people that identified in a different manner such as “Some Other Race” or “Two or more Races” (Figueroa-Lazu et al., 2022). A desire to identify with African ancestors or to reject the notions of the racial democracy or the adage *“hay que mejorar la raza”* (one has to improve/better the race) may be important factors in the formation of identity for some individuals. Therefore, individuals who identify as Afro-Puerto Rican may do so despite having a low percentage of African genetic ancestry or appearing phenotypically white by some cultural standard (Rivera, 2006).

The use of the 1KGP Puerto Rican population as a representative sample of Puerto Ricans that do not identify as Afro-descendant serves as a potential limitation of this study. These data are taken from a publicly available database that does not include information about how these individuals identify with respect to their African ancestry (Sudmant et al., 2015; The 1000 Genomes Consortium, 2015). It is therefore possible that some of the individuals present in the PUR population, if given the option, would choose to identify as Afro-Puerto Rican. Therefore, any statements made resulting from this research about possible genetic differences between Puerto Ricans that self-identify as Afro-Puerto Rican versus those that do not must consider the possibility of misclassification of some of the PUR individuals.

Additionally, Moreno-Estrada et al. (2013) have previously demonstrated distinct patterns of subcontinental African genetic ancestry in the Caribbean that differ between relatively short and long genetic ancestry tracts, indicating multiple migration pulses. As we did not perform tract-length analyses in this study, our subcontinental genetic ancestry results may be an average representation of differential patterns in shorter and longer tracts. Testing for the genetic presence of several distinct migration pulses, and resulting genetic ancestry patterns, for this population is therefore a possible direction for future work.

## 5. Conclusions

In conclusion, this study demonstrates that genetic approaches, when combined with ethnographically informed sampling (Benn Torres & Torres Colón, 2021; Winful et al., 2023) can reconstruct the demographic histories of self-identified African descendants in Puerto Rico. Our results suggest that while there are similarities with regard to general patterns of genetic ancestry among Afro-Caribbean communities, there is previously unrecognized regional heterogeneity within island sub-populations. Our findings are consistent with available historical sources, and also uncover novel information about the roles of African and European Ancestors in shaping the biocultural diversity of the modern Puerto Rican population.

## Acknowledgements

The authors would like to thank the study participants for trusting us with their DNA samples to conduct this research. We would also like to extend our deepest gratitude to Lisianette Laracuente and Alberto Galarza for their support and assistance with field sampling. We further thank Dr. Andrés Moreno Estrada for providing access to the bioinformatics resources of the Human Population and Evolutionary Genetics Laboratory at LANGEBIO-CINVESTAV. The authors acknowledge the Minnesota Supercomputing Institute (MSI) at University of Minnesota (UMN) for providing additional computing resources. This work was supported by funding from Vanderbilt University’s College of Arts and Sciences and The Office for Equity, Diversity and Inclusion, National Geographic Grant No.: NGS-68029R-20 awarded to J.B.T. Funds also provided by the National Science Foundation (NSF) SBE Postdoctoral Research Fellowship Award No. 1711982, and by new faculty startup funds from the UMN College of Liberal Arts, both awarded to M.A.N.C.

## Data Availability Statement

In efforts to maintain privacy, support autonomy, and address community concerns about future uses, all data remain under the control of the study community. Requests for project data may be made to the corresponding author (JBT) who will relay the request to the study community and help facilitate data sharing if approved.

## Author Contributions

**Maria Nieves-Colón:** Formal analysis (equal); methodology (equal); visualization (equal); data curation (equal); software (equal); writing - original draft (equal); writing - review and editing (equal). **Emma Ulrich:** Formal analysis (equal); methodology (equal); visualization (equal); data curation (equal); software (equal); writing - original draft (equal); writing - review and editing (equal). **Lijuan Chen:** Methodology (equal). **Gabriel Torres Colón:** Conceptualization (supporting); investigation (supporting); methodology (supporting); project administration (supporting); **Maricruz Rivera Clemente:** Conceptualization (supporting); investigation (supporting); project administration (supporting). **La Corporación Piñones Se Integra (COPI):** Conceptualization (supporting); investigation (supporting); project administration (supporting). **Jada Benn Torres:** Conceptualization (lead); data curation (lead); formal analysis (lead); funding acquisition (lead); investigation (lead); methodology (lead); project administration (lead); resources (lead); software (lead); supervision (lead); validation (lead); visualization (lead); writing - original draft (lead); writing - review and editing (lead).

## References

Adhikari, K., Chacón-Duque, J. C., Mendoza-Revilla, J., Fuentes-Guajardo, M., & Ruiz-Linares, A. (2017). The Genetic Diversity of the Americas. Annual Review of Genomics and Human Genetics, 18(1), 277–296. 10.1146/annurev-genom-083115-022331

Alexander, D. H., & Lange, K. (2011). Enhancements to the ADMIXTURE algorithm for individual ancestry estimation. BMC Bioinformatics, 12(1), 246. 10.1186/1471-2105-12-246

Alicea Rivera, A. (2017). El Tallit escondido: La presencia sefardita en Puerto Rico. Puerto Rico: Editorial Akelarre.

Altman, I. (2013). Marriage, Family, and Ethnicity in the Early Spanish Caribbean. The William and Mary Quarterly, 70(2), 225–250. 10.5309/willmaryquar.70.2.0225

Anderson, C. A., Pettersson, F. H., Clarke, G. M., Cardon, L. R., Morris, A. P., & Zondervan, K. T. (2010). Data quality control in genetic case-control association studies. Nature Protocols, 5(9), 1564–1573. 10.1038/nprot.2010.116

Ayala, C. J. (2007). Puerto Rico in the American century: A history since 1898. Chapel Hill: University of North Carolina Press.

Baralt, G. A. (2007). Slave revolts in Puerto Rico: Conspiracies and uprisings, 1795-1873. Princeton: Markus Wiener Publishers.

Behar, D. M., Yunusbayev, B., Metspalu, M., Metspalu, E., Rosset, S., Parik, J., … Villems, R. (2010). The genome-wide structure of the Jewish people. Nature, 466, 238–242. 10.1038/nature09103

Behr, A. A., Liu, K. Z., Liu-Fang, G., Nakka, P., & Ramachandran, S. (2016). pong: Fast analysis and visualization of latent clusters in population genetic data. Bioinformatics, 32(18), 2817–2823. 10.1093/bioinformatics/btw327

Benn Torres, J. (2014). Prospecting the past: Genetic perspectives on the extinction and survival of indigenous peoples of the Caribbean. New Genetics and Society, 33, 21–41. 10.1080/14636778.2013.873245

Benn Torres, J. (2018). ‘Reparational’ Genetics: Genomic Data and the Case for Reparations in the Caribbean. Genealogy, 2(1), 7. 10.3390/genealogy2010007

Benn Torres, J. (2020). Anthropological perspectives on genomic data, genetic ancestry, and race. Yearbook of Physical Anthropology, 171(S70), 74–86. 10.1002/ajpa.23979

Benn Torres, J., Kittles, R. A., & Stone, A. C. (2007). Mitochondrial and Y chromosome diversity in the English-speaking Caribbean. Annals of Human Genetics, 71, 782–790. 10.1111/j.1469-1809.2007.00380.x

Benn Torres, J., & Torres Colón, G. A. (2021). Genetic Ancestry: Our Stories, Our Pasts. Abingdon, Oxonlll; New York, NY: Routledge.

Bergström, A., McCarthy, S. A., Hui, R., Almarri, M. A., Ayub, Q., Danecek, P., … Tyler-Smith, C. (2020). Insights into human genetic variation and population history from 929 diverse genomes. Science, 367(6484), eaay5012. 10.1126/science.aay5012

Bertoni, B., Budowle, B., Sans, M., Barton, S. A., & Chakraborty, R. (2003). Admixture in Hispanics: Distribution of ancestral population contributions in the Continental United States. Hum Biol, 75, 1–11. 10.1353/hub.2003.0016

Browning, S. R., & Browning, B. L. (2007). Rapid and accurate haplotype phasing and missing-data inference for whole-genome association studies by use of localized haplotype clustering. American Journal of Human Genetics, 81(5), 1084–1097. 10.1086/521987

Browning, S. R., Grinde, K., Plantinga, A., Gogarten, S. M., Stilp, A. M., Kaplan, R. C., … Laurie, C. C. (2016). Local Ancestry Inference in a Large US-Based Hispanic/Latino Study: Hispanic Community Health Study/Study of Latinos (HCHS/SOL). G3: Genes|Genomes|Genetics, 6(6), 1525–1534. 10.1534/g3.116.028779

Bryc, K., Auton, A., Nelson, M. R., Oksenberg, J. R., Hauser, S. L., Williams, S., … Bustamante, C. D. (2010). Genome-wide patterns of population structure and admixture in West Africans and African Americans. Proceedings of the National Academy of Sciences, 107(2), 786–791. 10.1073/pnas.0909559107

Bukhari, A., Luis, J. R., Alfonso-Sanchez, M. A., Garcia-Bertrand, R., & Herrera, R. J. (2017). Taino and African maternal heritage in the Greater Antilles. Gene, 637, 33–40. 10.1016/j.gene.2017.09.004

Central Intelligence, A. (2021, June 9). Puerto Rico—The World Factbook. Retrieved June 23, 2021, from The World Factbook website: https://www.cia.gov/the-world-factbook/countries/puerto-rico/#people-and-society

Chacón-Duque, J.-C., Adhikari, K., Fuentes-Guajardo, M., Mendoza-Revilla, J., Acuña-Alonzo, V., Barquera, R., … Ruiz-Linares, A. (2018). Latin Americans show wide-spread Converso ancestry and imprint of local Native ancestry on physical appearance. Nature Communications, 9(1), 5388. 10.1038/s41467-018-07748-z

Chang, C. C., Chow, C. C., Tellier, L. C., Vattikuti, S., Purcell, S. M., & Lee, J. J. (2015). Second-generation PLINK: Rising to the challenge of larger and richer datasets. GigaScience, 4(1), s13742–015-0047–0048. 10.1186/s13742-015-0047-8

Chiang, C. W. K., Marcus, J. H., Sidore, C., Biddanda, A., Al-Asadi, H., Zoledziewska, M., … Novembre, J. (2018). Genomic history of the Sardinian population. Nature Genetics, 50(10), 1426–1434. 10.1038/s41588-018-0215-8

Chinea, J. (1996). Race, Colonial Exploitation and West Indian Immigration in Nineteenth-Century Puerto Rico, 1800-1850. The Americas, 52(4), 495–519. 10.2307/1008475

Chinea, J. (1997). A Quest for Freedom: The Immigration of Maritime Maroons into Puerto RIco, 1656-1800. The Journal of Caribbean History, 31(1/2), 51–87.

Chinea, J. (2016). Slavery and Child Trafficking in Puerto Rico at the Closing of the African Slave Trade: The Young Captives of the Slaver Majesty, 1859-1865. Revista Brasileira Do Caribe, 17(32), 41.

Cifre de Loubriel, E. (1964). La inmigración a Puerto Rico durante el Siglo XIX. Instituto de Cultura Puertorriqueña.

Cox, T. F., & Cox, M. A. A. (1994). Multidimensional Scaling. Taylor & Francis.

Curet, L. A. (2005). Caribbean paleodemography: Population, culture history, and sociopolitical processes in ancient Puerto Rico. Tuscaloosa, Ala: University of Alabama Press.

Demby, G., & Meraji, S. M. (2020, April 24). Puerto Rico, Island Of Racial Harmony? In Code Switch. Retrieved from https://www.npr.org/transcripts/842832544

Díaz-Zabala, H. J., Nieves-Colón, M. A., & Martínez-Cruzado, J. C. (2017). A Mainly Circum-Mediterranean Origin for West Eurasian and North African mtDNAs in Puerto Rico with Strong Contributions from the Canary Islands and West Africa. Human Biology, 89(2), 125. 10.13110/humanbiology.89.2.04

Elhaik, E., Greenspan, E., Staats, S., Krahn, T., Tyler-Smith, C., Xue, Y., … Genographic, C. (2013). The GenoChip: A new tool for genetic anthropology. Genome Biology and Evolution, 5, 1021–1031. 10.1093/gbe/evt066

Eltis, D. (2009). A Brief Overview of the Trans-Atlantic Slave Trade,” Voyages: The Trans-Atlantic Slave Trade Database. Retrieved from A Brief Overview of the Trans-Atlantic Slave Trade,’ Slave Voyages: The Trans-Atlantic Slave Trade Database website: https://www.slavevoyages.org/voyage/about

Fernandes, D. M., Sirak, K. A., Ringbauer, H., Sedig, J., Rohland, N., Cheronet, O., … Reich, D. (2021). A genetic history of the pre-contact Caribbean. Nature, 590(7844), 103–110. 10.1038/s41586-020-03053-2

Fernandez Cobo, M., Jobes, D., Yanagihara, R., Nerurkar R, Y. Y., Ryschkewitsch, C., & Stoner, G. (2001). Reconstructing population history using JC virus: Amerinds, Spanish, and Africans in the ancestry of modern Puerto Ricans. Human Biology, 73, 385–402. 10.1353/hub.2001.0032

Figueroa, L. A. (2005). Sugar, Slavery, and Freedom in Nineteenth-Century Puerto Rico. Chapel Hill: The University of North Carolina Press.

Figueroa-Lazu, D., Hinojosa, J., & Bonilla, Y. (2022). Puerto Rico’s 2020 Race/Ethnicity Decennial Analysis. New York, NY: Center for Puerto Rican Studies, Hunter College. Retrieved from Center for Puerto Rican Studies, Hunter College website: https://centropr.hunter.cuny.edu/app/uploads/2023/02/022223-Puerto-Ricos-2020-Race_Ethnicity-Decennial-Analysis.pdf

Flores-Bello, A., Bauduer, F., Salaberria, J., Oyharçabal, B., Calafell, F., Bertranpetit, J., … Comas, D. (2021). Genetic origins, singularity, and heterogeneity of Basques. Current Biology, 31(10), 2167–2177.e4. 10.1016/j.cub.2021.03.010

Fortes-Lima, C., Gessain, A., Ruiz-Linares, A., Bortolini, M.-C., Migot-Nabias, F., Bellis, G., … Dugoujon, J.-M. (2017). Genome-wide Ancestry and Demographic History of African-Descendant Maroon Communities from French Guiana and Suriname. The American Journal of Human Genetics, 101(5), 725–736. 10.1016/j.ajhg.2017.09.021

García Leduc, J. M. (2002). Apuntes para una historia breve de Puerto Rico: Desde la prehistoria hasta 1898. San Juan, P.R: Editorial Isla Negra. Retrieved from https://catalog.hathitrust.org/Record/101250648

Godreau, I. P. (2015). Scripts of Blackness: Race, Cultural Nationalism, and US Colonialism in Puerto Rico. University of Illinois Press.

Godreau, I. P., & Bonilla, Y. (2021). Nonsovereign Racecraft: How Colonialism, Debt, and Disaster are Transforming Puerto Rican Racial Subjectivities. American Anthropologist, 124(3), 509–525. 10.1111/aman.13601

Gouveia, M. H., Borda, V., Leal, T. P., Moreira, R. G., Bergen, A. W., Kehdy, F. S. G., … Tarazona-Santos, E. (2020). Origins, Admixture Dynamics, and Homogenization of the African Gene Pool in the Americas. Molecular Biology and Evolution, 37(6), 1647–1656. 10.1093/molbev/msaa033

Gower, J. C. (1966). Some distance properties of latent root and vector methods used in multivariate analysis. Biometrika, 53(3–4), 325–338. 10.1093/biomet/53.3-4.325

Gurdasani, D., Carstensen, T., Tekola-Ayele, F., Pagani, L., Tachmazidou, I., Hatzikotoulas, K., … Sandhu, M. S. (2015). The African Genome Variation Project shapes medical genetics in Africa. Nature, 517(7534), 327–332. 10.1038/nature13997

Hall, G. M. (2005). Slavery and African Ethnicities in the Americas: Restoring the Links. Chapel Hill: The University of North Carolina Press. 10.5149/9780807876862_hall

Hernández, C. L., Pita, G., Cavadas, B., López, S., Sánchez-Martínez, L. J., Dugoujon, J.-M., … Calderón, R. (2020). Human Genomic Diversity Where the Mediterranean Joins the Atlantic. Molecular Biology and Evolution, 37(4), 1041–1055. 10.1093/molbev/msz288

Hinojosa, J. (2018). Two sides of the coin of puerto rican migration: Depopulation in puerto rico and the redefinition of the diaspora. CENTRO: Journal of the Center for Puerto Rican Studies, 30(3), 230–253. Gale Academic OneFile. Retrieved from Gale Academic OneFile.

Hofman, C. L., Borck, L., Laffoon, J. E., Slayton, E. R., Scott, R. B., Breukel, T. W., … Hoogland, M. L. P. (2020). Island networks: Transformations of inter-community social relationships in the Lesser Antilles at the advent of European colonialism. The Journal of Island and Coastal Archaeology, 0(0), 1–27. 10.1080/15564894.2020.1748770

Jackson, F. L. C. (2021). So many Nigerians: Why is Nigeria overrepresented as the ancestral genetic homeland of Legacy African North Americans? The American Journal of Human Genetics, 108, 1–7. 10.1016/j.ajhg.2020.10.010

Kagan, R. L., & Morgan, P. D. (Eds.). (2009). Atlantic Diasporas: Jews, Conversos and Crypto-Jews in the Age of Mercantilism. Baltimore: Johns Hopkins University Press.

Knight, F. W. (Ed.). (2003). General History of the Caribbean: Volume III: The slave societies of the Caribbean. New York: Palgrave Macmillan US. 10.1007/978-1-349-73770-3

Laffoon, J. E., Rodríguez Ramos, R., Chanlatte Baik, L., Narganes Storde, Y., Rodríguez Lopez, M., Davies, G. R., & Hofman, C. L. (2014). Long-distance exchange in the precolonial Circum-Caribbean: A multi-isotope study of animal tooth pendants from Puerto Rico. Journal of Anthropological Archaeology, 35, 220–233. 10.1016/j.jaa.2014.06.004

Laó-Montes, A. (2018). Afro-Boricua Agency. ReVista (Cambridge), 17(2), 12–15.

Las Casas, B. (2004). A Short Account of the Destruction of the Indies (Vol. 2). London, United Kingdom: Penguin Books Limited.

Lloréns, H., García-Quijano, C. G., & Godreau, I. P. (2017). Racismo en Puerto Rico: Surveying Perceptions of Racism. CENTRO: Journal of the Center for Puerto Rican Studies, 30.

Maples, B. K., Gravel, S., Kenny, E. E., & Bustamante, C. D. (2013). RFMix: A discriminative modeling approach for rapid and robust local-ancestry inference. American Journal of Human Genetics, 93(2), 278–288. 10.1016/j.ajhg.2013.06.020

Marcus, J. H., Posth, C., Ringbauer, H., Lai, L., Skeates, R., Sidore, C., … Novembre, J. (2020). Genetic history from the Middle Neolithic to present on the Mediterranean island of Sardinia. Nature Communications, 11(1), 939. 10.1038/s41467-020-14523-6

Martínez-Cruzado, J. C., Toro-Labrador, G., Viera-Vera, J., Rivera-Vega, M. Y., Startek, J., Latorre-Esteves, M., … Valencia-Rivera, P. (2005). Reconstructing the population history of Puerto Rico by means of mtDNA phylogeographic analysis. American Journal of Physical Anthropology, 128(1), 131–155. 10.1002/ajpa.20108

Mayo Santana, R., & Negrón Portillo, M. (2007). La esclavitud menor: La esclavitud en los municipios del interior de Puerto Rico en el siglo XIX. San Juan, P.R: CIS, Centro de Investiagciones Sociales, Universidad de Puerto Rico. Retrieved from http://bibliotecavirtual.clacso.org.ar/Puerto_Rico/cis-uprrp/20120806105028/entero.pdf

Mendisco, F., Pemonge, M., Romon, T., Lafleur, G., Richard, G., Courtaud, P., & Deguilloux, M. (2019). Tracing the genetic legacy in the French Caribbean islands: A study of mitochondrial and Y-chromosome lineages in the Guadeloupe archipelago. American Journal of Physical Anthropology, 170(4), 507–518. 10.1002/ajpa.23931

Montinaro, F., Busby, G. B. J., Pascali, V. L., Myers, S., Hellenthal, G., & Capelli, C. (2015). Unravelling the hidden ancestry of American admixed populations. Nature Communications, 6(1), 6596. 10.1038/ncomms7596

Moore, D. S., McCabe, G. P., & Craig, B. A. (2007). Introduction to the Practice of Statistics: W/Student CD (6th edition). Houndmills: W. H. Freeman.

Moreno-Estrada, A., Gravel, S., Zakharia, F., McCauley, J. L., Byrnes, J. K., Gignoux, C. R., … Bustamante, C. D. (2013). Reconstructing the population genetic history of the Caribbean. PLoS Genetics, 9, e1003925. 10.1371/journal.pgen.1003925

Moscoso, F. (1995). Formas de resistencia de los esclavos en Puerto Rico. Siglos XVI-XVIII. América Negra, 10, 31–48.

Murray, T., Beaty, T. H., Mathias, R. A., Rafaels, N., Grant, A. V., Faruque, M. U., … Barnes, K. C. (2010). African and non-African admixture components in African Americans and an African Caribbean population. Genetic Epidemiology, 34(6), 561–568. 10.1002/gepi.20512

Nägele, K., Posth, C., Orbegozo, M. I., Armas, Y. C. de, Godoy, S. T. H., Herrera, U. M. G., … Schroeder, H. (2020). Genomic insights into the early peopling of the Caribbean. Science, 369(6502), 456–460. 10.1126/science.aba8697

Nieves-Colón, M. A. (2022). Anthropological genetic insights on Caribbean population history. *Evolutionary Anthropology: Issues*, News, and Reviews, 31(3), 118–137. 10.1002/evan.21935

Nieves-Colón, M. A., Pestle, W. J., Reynolds, A. W., Llamas, B., de la Fuente, C., Fowler, K., … Stone, A. C. (2020). Ancient DNA Reconstructs the Genetic Legacies of Precontact Puerto Rico Communities. Molecular Biology and Evolution, 37(3), 611–626. 10.1093/molbev/msz267

Nistal-Moret, B. (2004). Esclavos prófugos y cimarrones: Puerto Rico, 1770-1870. San Juan, P.R: Editorial de la Universidad de Puerto Rico.

Ostrer, H., & Skorecki, K. (2013). The population genetics of the Jewish people. Human Genetics, 132(2), 119–127. 10.1007/s00439-012-1235-6

Patin, E., Lopez, M., Grollemund, R., Verdu, P., Harmant, C., Quach, H., … Froment, A. (2017). Dispersals and genetic adaptation of Bantu-speaking populations in Africa and North America. Science, 356(6337), 543–546. 10.1126/science.aal1988

Patterson, N., Price, A. L., & Reich, D. (2006). Population Structure and Eigenanalysis. PLoS Genetics, 2(12), e190. 10.1371/journal.pgen.0020190

Pestle, W. J., Perez, E. M., & Koski-Karell, D. (2023). Reconsidering the lives of the earliest Puerto Ricans: Mortuary Archaeology and bioarchaeology of the Ortiz site. PLOS ONE, 18(4), e0284291. 10.1371/journal.pone.0284291

Picó, F. (2008). Historia general de Puerto Rico (4a *ed.)*. San Juan, P.R: Ediciones Huracán. Retrieved from https://www.digitaliapublishing.com/a/28728/historia-general-depuerto-rico--4a-ed.-

Price, A. L., Patterson, N. J., Plenge, R. M., Weinblatt, M. E., Shadick, N. A., & Reich, D. (2006). Principal components analysis corrects for stratification in genome-wide association studies. Nature Genetics, 38(8), 904–909. 10.1038/ng1847

R Core Team. (2020). R: A Language and Environment for Statistical Computing. Vienna, Austria: R Foundation for Statistical Computing. Retrieved from https://www.R-project.org/

Rivera, M. Q. (2006). From Trigueñita to Afro-Puerto Rican: Intersections of the Racialized, Gendered, and Sexualized Body in Puerto Rico and the U.S. Mainland. Meridians, 7(1), 162–182. 10.2979/MER.2006.7.1.162

Rivera-Rideau, P. R. (2013). From Carolina to Loíza: Race, place and Puerto Rican racial democracy. Identities, 20(5), 616–632. 10.1080/1070289X.2013.842476

Rodríguez Mendoza, F. (2004). La emigración del noroeste de Tenerife a América durante 1750-1830. Santa Cruz de Tenerife, Spain: Servicio de Publicaciones, Universidad de La Laguna. Retrieved from http://riull.ull.es/xmlui/handle/915/9909

Rodríguez Ramos, R., Rodríguez López, M., & Pestle, W. J. (2023). Revision of the cultural chronology of precolonial Puerto Rico: A Bayesian approach. PLOS ONE, 18(2), e0282052. 10.1371/journal.pone.0282052

Rodríguez-Rodríguez, J. E., Ioannidis, A. G., Medina-Muñoz, S. G., Barberena-Jonas, C., Blanco-Portillo, J., Quinto-Cortés, C. D., & Moreno-Estrada, A. (2022). The genetic legacy of the Manila galleon trade in Mexico. Philosophical Transactions of the Royal Society of London. Series B, Biological Sciences, 377(1852), 20200419. 10.1098/rstb.2020.0419

Rodríguez-Silva, I. (2012). Silencing race: Disentangling blackness, colonialism, and national identities in Puerto Rico. Springer.

Santana Pérez, J. M. (2008). Isleños en Cuba y Puerto Rico (del siglo XVII a mediados del XIX). Cuadernos Americanos: Nueva Epoca, 4(126), 173–192.

Scarano, F. A. (1981). Inmigración y clases sociales en el Puerto Rico del siglo XIX. Río Piedras, P.R.: Ediciones Huracán.

Schroeder, H., Ávila-Arcos, M. C., Malaspinas, A.-S., Poznik, G. D., Sandoval-Velasco, M., Carpenter, M. L., … Gilbert, M. T. P. (2015). Genome-wide ancestry of 17th-century enslaved Africans from the Caribbean. Proceedings of the National Academy of Sciences of the United States of America, 112(12), 3669–3673. 10.1073/pnas.1421784112

Schwartz, S. B. (1997). Spaniards, “Pardos”, and the Missing Mestizos: Identities and Racial Categories in the Early Hispanic Caribbean. NWIG: New West Indian Guide / Nieuwe West-Indische Gids, 71(1/2), 5–19.

Shi, J., O’Connell, J., Hicks, B., Wang, W., Bryc, K., Brady, J. J., … Shringarpure, S. (2023, July 28). GWAS of cataract in Puerto Ricans identifies a novel large-effect variant in ITGA6. 10.1101/2023.07.25.23293173

Siegel, P. E. (2005). Ancient Borinquen archaeology and ethnohistory of native Puerto Rico. Tuscaloosa: University of Alabama Press.

Stark, D. M. (2009). A New Look at the African Slave Trade in Puerto Rico Through the Use of Parish Registers: 1660–1815. Slavery & Abolition, 30, 491–520. 10.1080/01440390903245083

Stefflova, K., Dulik, M. C., Barnholtz-Sloan, J. S., Pai, A. A., Walker, A. H., & Rebbeck, T. R. (2011). Dissecting the within-Africa ancestry of populations of African descent in the Americas. PLOS ONE, 6, e14495. 10.1371/journal.pone.0014495

Sudmant, P. H., Rausch, T., Gardner, E. J., Handsaker, R. E., Abyzov, A., Huddleston, J., … Korbel, J. O. (2015). An integrated map of structural variation in 2,504 human genomes. Nature, 526(7571), 75–81. 10.1038/nature15394

The 1000 Genomes Consortium. (2015). A global reference for human genetic variation. Nature, 526(7571), 68–74. 10.1038/nature15393

Vargas-Ramos, C. (2005). BLACK, TRIGUEÑO, WHITE …? Shifting Racial Identification among Puerto Ricans. Du Bois Review: Social Science Research on Race, 2(2), 267–285. 10.1017/S1742058X05050186

Via, M., Gignoux, C. R., Roth, L. A., Fejerman, L., Galanter, J., Choudhry, S., … Martínez-Cruzado, J. C. (2011). History Shaped the Geographic Distribution of Genomic Admixture on the Island of Puerto Rico. PLOS ONE, 6(1), e16513. 10.1371/journal.pone.0016513

Vilar, M. G., Melendez, C., Sanders, A. B., Walia, A., Gaieski, J. B., Owings, A. C., & Schurr, T. G. (2014). Genetic diversity in Puerto Rico and its implications for the peopling of the island and the West Indies. American Journal of Physical Anthropology, 155, 352–386. 10.1002/ajpa.22569

Wheat, D. (2016). Atlantic Africa and the Spanish Caribbean, 1570-1640. Chapel Hill: The University of North Carolina Press. 10.5149/9781469623801_wheat

Wickham, H. (2016). ggplot2: Elegant Graphics for Data Analysis. New York: Springer-Verlag. Retrieved from https://link.springer.com/book/10.1007/978-3-319-24277-4

Winful, T., Mccormack, K., Mueller, E., Chen, L., La Corporación Piñones Se Integra (COPI), Rivera Clemente, M., & Benn Torres, J. (2023). Exploring the legacy of African and Indigenous Caribbean admixture in Puerto Rico. American Journal of Biological Anthropology. 10.1002/ajpa.24814

Wojcik, G. L., Graff, M., Nishimura, K. K., Tao, R., Haessler, J., Gignoux, C. R., … Carlson, C. S. (2019). Genetic analyses of diverse populations improves discovery for complex traits. Nature, 570(7762), 514–518. 10.1038/s41586-019-1310-4

