## Supplementary material for "Genetic ancestry in Puerto Rican Afro-descendants illustrates diverse histories of African diasporic populations": Figure S1

* Co-first authors
[†] Corresponding author

Supporting Tables S1-S5 (in excel file SuppInfo_Tables.xlsx)
Supporting Figures S1-S12 (in Supporting Information file)
This Supporting Information file contains Supporting Figures S1-S12.

*Six populations from the African Genome Variation Project (Gurdasani et al., 2015) panel were individually QC'd according to the same process outlined below and then merged using PLINK 1.9. After QC, 518 individuals remained across these AGVP populations. When these populations were merged into the full African subcontinental panel, the overlap was 526,960 SNPs.*

| | **Afro-Puerto Rican sample individuals** (711,059 SNPs, N=30) | **Population Architecture Using Genomics and Epidemiology (Puno)** (1,204,593 SNPs, N=100) | **Patin et al., 2017 (Bantu-speaking populations)** (690,739 SNPs, N=1,202) | **1000 Genomes Project autosomal SNP data (African Panel)** (77,818,345 SNPs, N=398) | **Behar et al., 2010 (Jewish Diaspora)** (555,737 SNPs, N=87) | **Human Genome Diversity Project (European Panel)** (535,114 SNPs, N=140) | **Hernández et al., 2020 (Western Mediterranean)** (2,372,784 SNPs, N=104) | **1000 Genomes Project (European Panel)** (84,358,431 SNPs, N=778) |
|---|---|---|---|---|---|---|---|---|
| **QC Filter 1:** Restrict to autosomes | 30,176 SNPs removed (680,883 SNPs, N=30) | 0 SNPs removed (1,204,593 SNPs, N=100) | 0 SNPs removed (690,739 SNPs, N=1,202) | 0 SNPs removed (77,818,345 SNPs, N=398) | 13,374 SNPs removed (542,363 SNPs, N=87) | 0 SNPs removed (535,114 SNPs, N=140) | 60,204 SNPs removed (2,312,580 SNPs, N=104) | 3,502,709 SNPs removed (80,855,722 SNPs, N=778) |
| **QC Filter 2:** Restrict to SNP-only variants | 42 SNPs removed (680,841 SNPs, N=30) | 17,051 SNPs removed (1,187,542, N=100) | 0 SNPs removed (690,739 SNPs, N=1,202) | 0 SNPs removed (77,818,345 SNPs, N=398) | 0 SNPs removed (542,363 SNPs, N=87) | 0 SNPs removed (535,114 SNPs, N=140) | 0 SNPs removed (2,312,580 SNPs, N=104) | 3,037,377 SNPs removed (77,818,345 SNPs, N=778) |
| **QC Filter 3:** Remove duplicate variants | 347 SNPs removed (680.494 SNPs, N=30) | 2 SNPs removed (1,187,540 SNPs, N=100) | 0 SNPs removed (690,739 SNPs, N=1,202) | 0 SNPs removed (77,818,345 SNPs, N=398) | 0 SNPs removed (542,363 SNPs, N=87) | 0 SNPs removed (535,114 SNPs, N=140) | 0 SNPs removed (2,312,580 SNPs, N=104) | 626 SNPs removed (77,817,719 SNPs, N=778) |
| **QC Filter 4:** Filter by --mind 0.10 and --geno 0.05 | 13,453 SNPs removed 0 individuals removed (667,041 SNPs, N=30) | 1,360 SNPs removed 0 individuals removed (1,186,180 SNPs, N=100) | 195 SNPs removed 0 individuals removed (690,544 SNPs, N=1,202) | 0 SNPs removed 0 individuals removed (77,818,345 SNPs, N=398) | 481 SNPs removed 0 individuals removed (541,882 SNPs, N=87) | 469 SNPs removed 0 individuals removed (534,645 SNPs, N=140) | 41,256 SNPs removed 0 individuals removed (2,271,324 SNPs, N=104) | 0 SNPs removed 0 individuals removed (77,817,719 SNPs, N=778) |
| **QC Filter 5:** Filter by --hwe 1e-5 | 36 SNPs removed (667,005 SNPs, N=30) | 3 SNPs removed (1,186,177 SNPs, N=100) | 2,226 SNPs removed (688,318 SNPs, N=1,202) | 83,207 SNPs removed (77,735,138 SNPs, N=398) | 23 SNPs removed (541,859 SNPs, N=87) | 7 SNPs removed (534,638 SNPs, N=140) | 165 SNPs removed (2,271,159 SNPs, N=104) | 933,763 SNPs removed (76,883,956 SNPs, N=778) |
| **QC Filter 6:** Remove SNPs marked as ambiguous by snpflip | 4,336 SNPs removed **(662,669 SNPs, N=30)** | 0 SNPs removed **(1,186,177 SNPs, N=100)** | 637 SNPs removed **(687,681 SNPs, N=1,202)** | SNPs removed **(77,735,138 SNPs, N=398)** | 10 SNPs removed **(541,849 SNPs, N=87)** | 0 SNPs removed **(534,638 SNPs, N=140)** | 77,792 SNPs removed **(2,193,367 SNPs, N=104)** | 11,996,586 SNPs removed **(64,887,370 SNPs, N=778)** |

**Figure S1.** Quality-control (QC) filtering per each PLINK-formatted dataset used in this study.

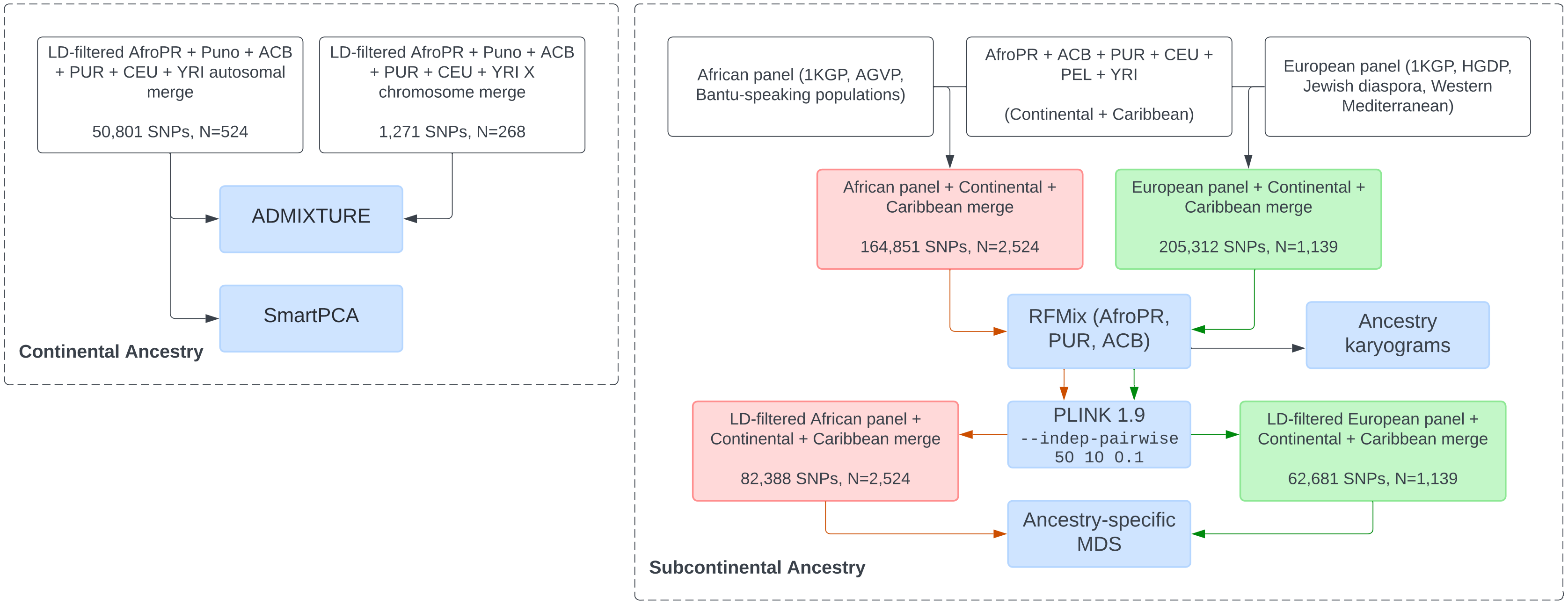


**Figure S2.** Workflow of analyses and merged datasets used in this study. Analyses of continental ancestry (left) include ADMIXTURE and SmartPCA. Analyses of subcontinental African and European ancestry (right) include RFMix, ancestry-specific MDS as outlined by Browning et al. 2016, and the production of ancestry karyograms. Three-letter population codes (ACB, PUR, CEU, YRI, PEL) correspond to 1000 Genomes Project populations.

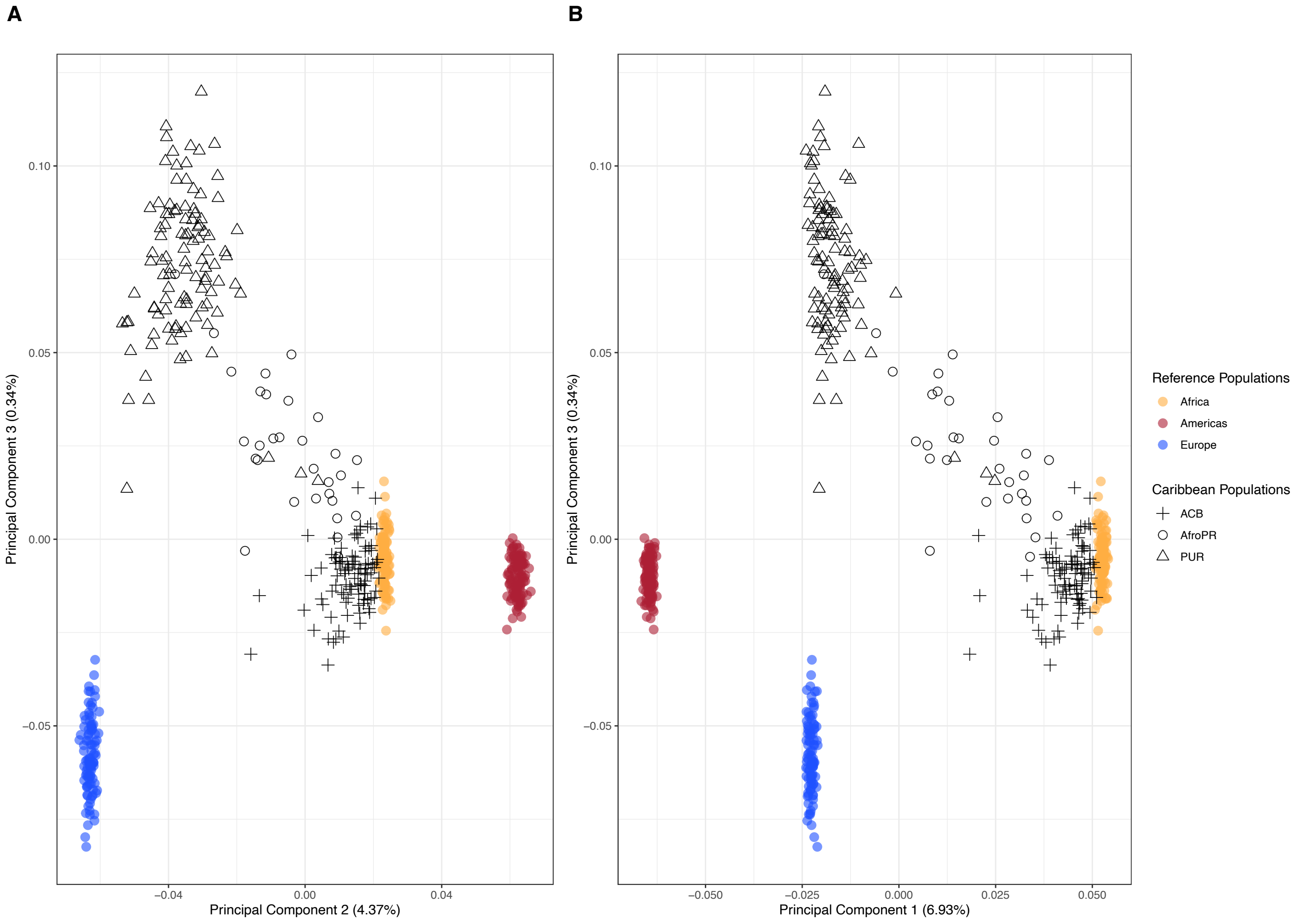


**Figure S3.** (A) Principal component axes 2 vs. 3 and (B) principal component axes 1 vs. 3 of the SmartPCA analysis of AfroPR sample individuals, two 1000 Genomes Project Caribbean populations, and three 1000 Genomes Project continental references (colored circles). Study population individuals are represented by open circles, PUR individuals by open triangles, and ACB individuals by crosses.

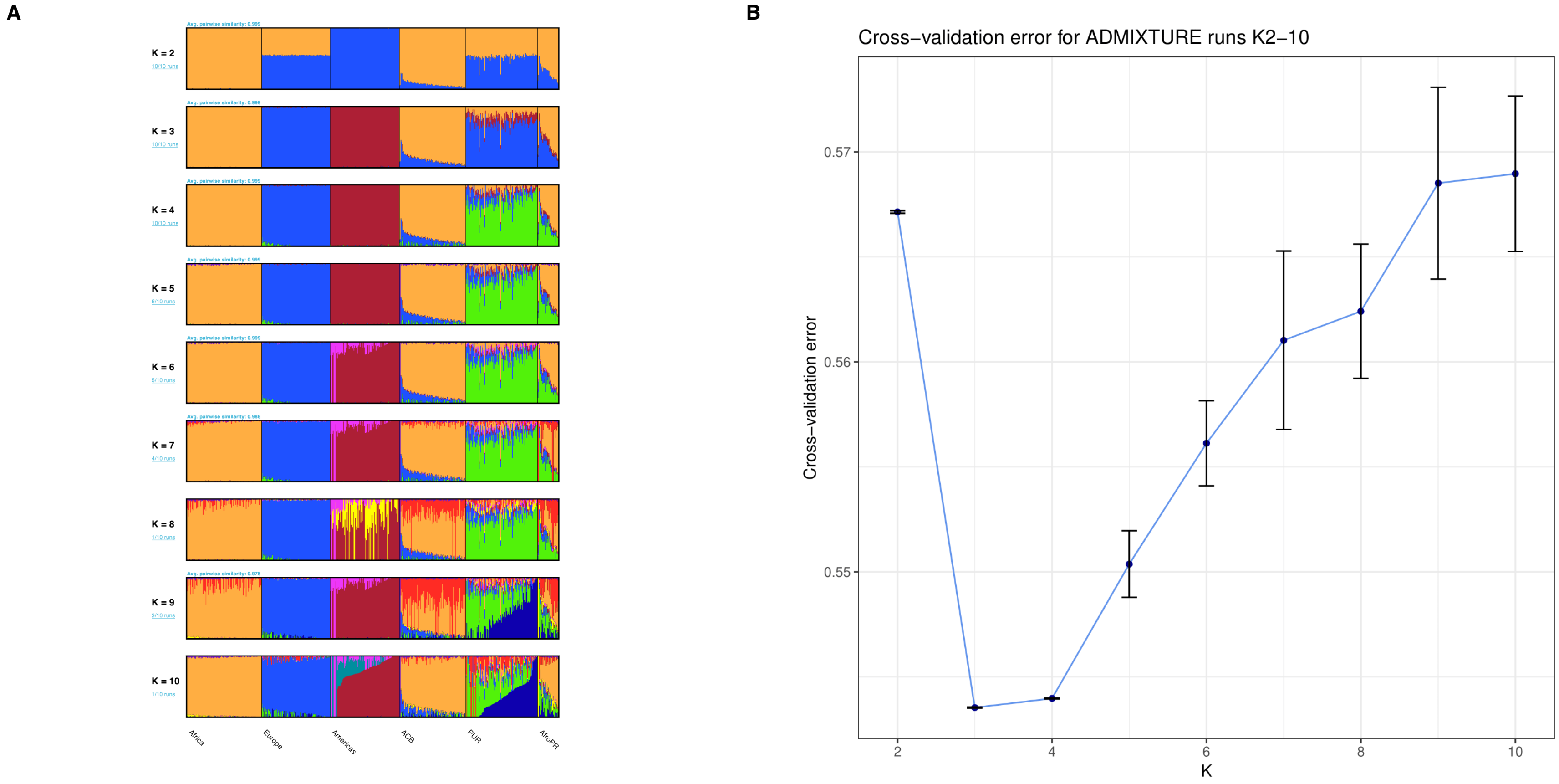


**Figure S4: ADMIXTURE analysis and cross-validation error of the study population and comparative regional populations (autosomal).** (A) ADMIXTURE clustering results for models assuming K=2 through K=10 clusters. Thin vertical bars represent individuals, with thin black bars separating populations. Continental reference populations and additional Caribbean populations are selected from the 1000 Genomes Project. (B) Cross-validation error computed for the ADMIXTURE K=2 through K=10 models. The model assuming K=3 is associated with the lowest cross-validation error.

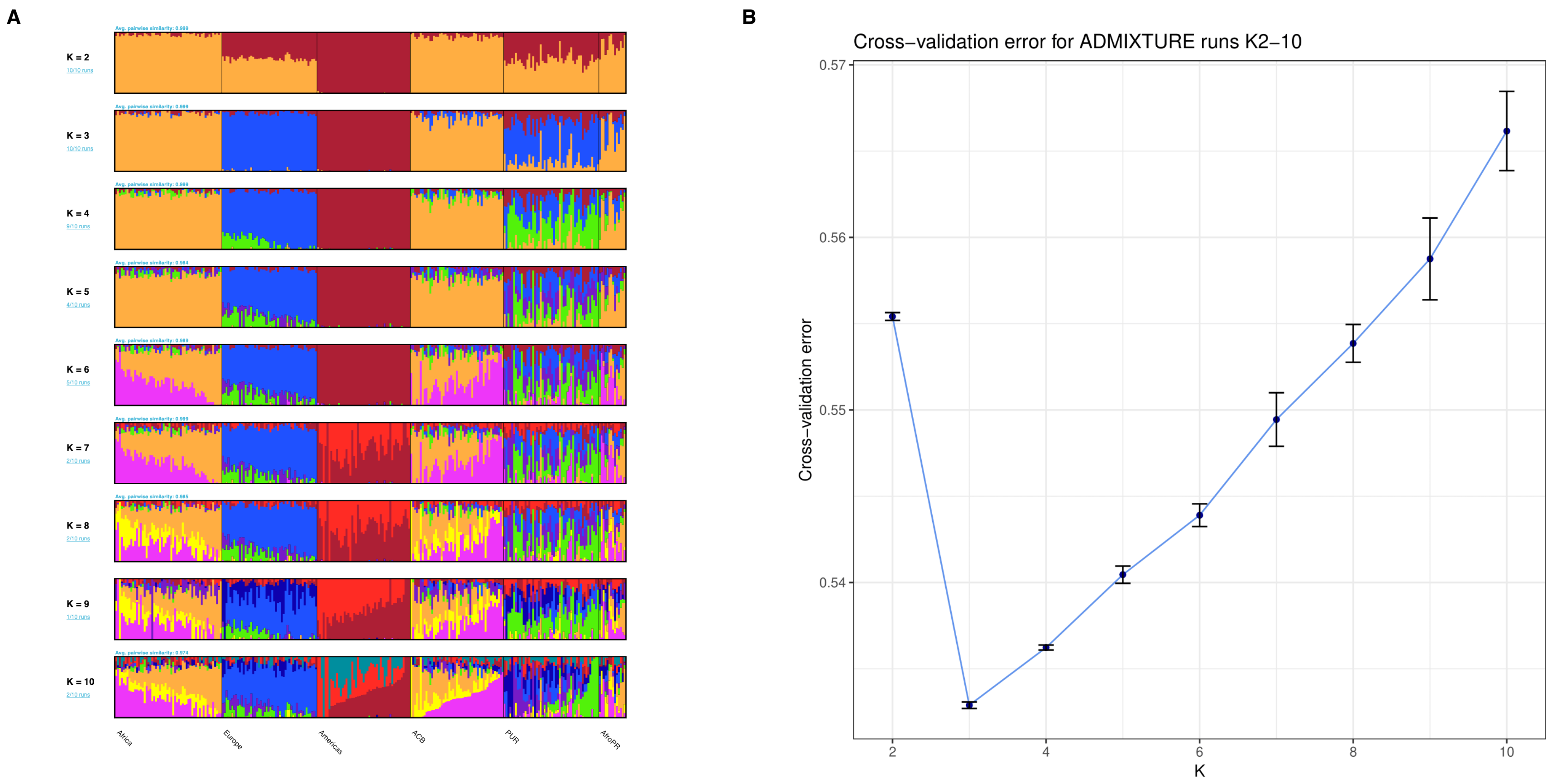


**Figure S5: ADMIXTURE analysis and cross-validation error of chromosomal females from the study population and comparative regional populations (X chromosome).** (A) ADMIXTURE clustering results performed on X chromosome data for models assuming K=2 through K=10 clusters. Thin vertical bars represent individuals, with thin black bars separating populations. All individuals included in this analysis are chromosomal females (two X chromosomes). Continental reference populations and additional Caribbean populations are selected from the 1000 Genomes Project. (B) Cross-validation error computed for the X chromosome ADMIXTURE K=2 through K=10 models. The model assuming K=3 is associated with the lowest cross-validation error.

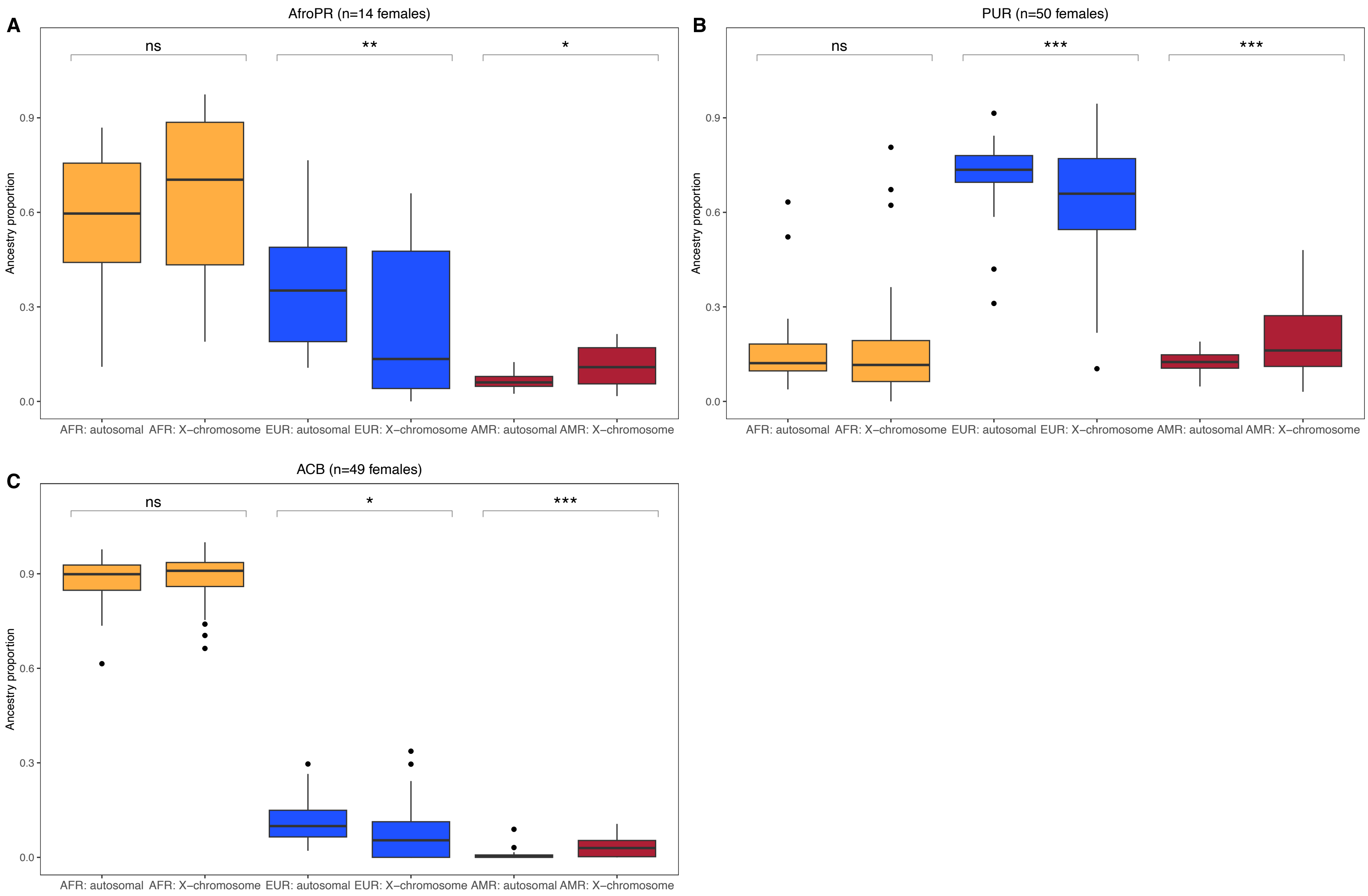


**Figure S6: Comparison of population structure on the X chromosome and autosomal chromosomes for the study population**

**and comparative regional populations.** (A) Boxplots with associated Wilcoxon-rank sum calculated significance values comparing African, European, and American ancestry proportions on the X chromosome and all autosomal chromosomes for chromosomal females from the study population. No significant difference was found for the African ancestry proportion, but significance corresponding to $0.001 \leq p \leq 0.01$ was found for the European ancestry proportion and to $0.01 \leq p \leq 0.05$ for the American ancestry proportion. (B) Boxplots with associated Wilcoxon-rank sum calculated significance values comparing African, European, and American ancestry proportions on the X chromosome and all autosomal chromosomes for chromosomal females from the 1000 Genomes Project PUR population. No significant difference was found for the African ancestry proportion, but significance corresponding to $p \leq 0.001$ was found for the European and American ancestry proportions. (C) Boxplots with associated Wilcoxon-rank sum calculated significance values comparing African, European, and American ancestry proportions on the X chromosome and all autosomal chromosomes for chromosomal females from the 1000 Genomes Project ACB population. No significant difference was found for the African ancestry proportion, but significance corresponding to $0.01 \leq p \leq 0.05$ was found for the European ancestry proportion and to $p \leq 0.001$ for the American ancestry proportion.

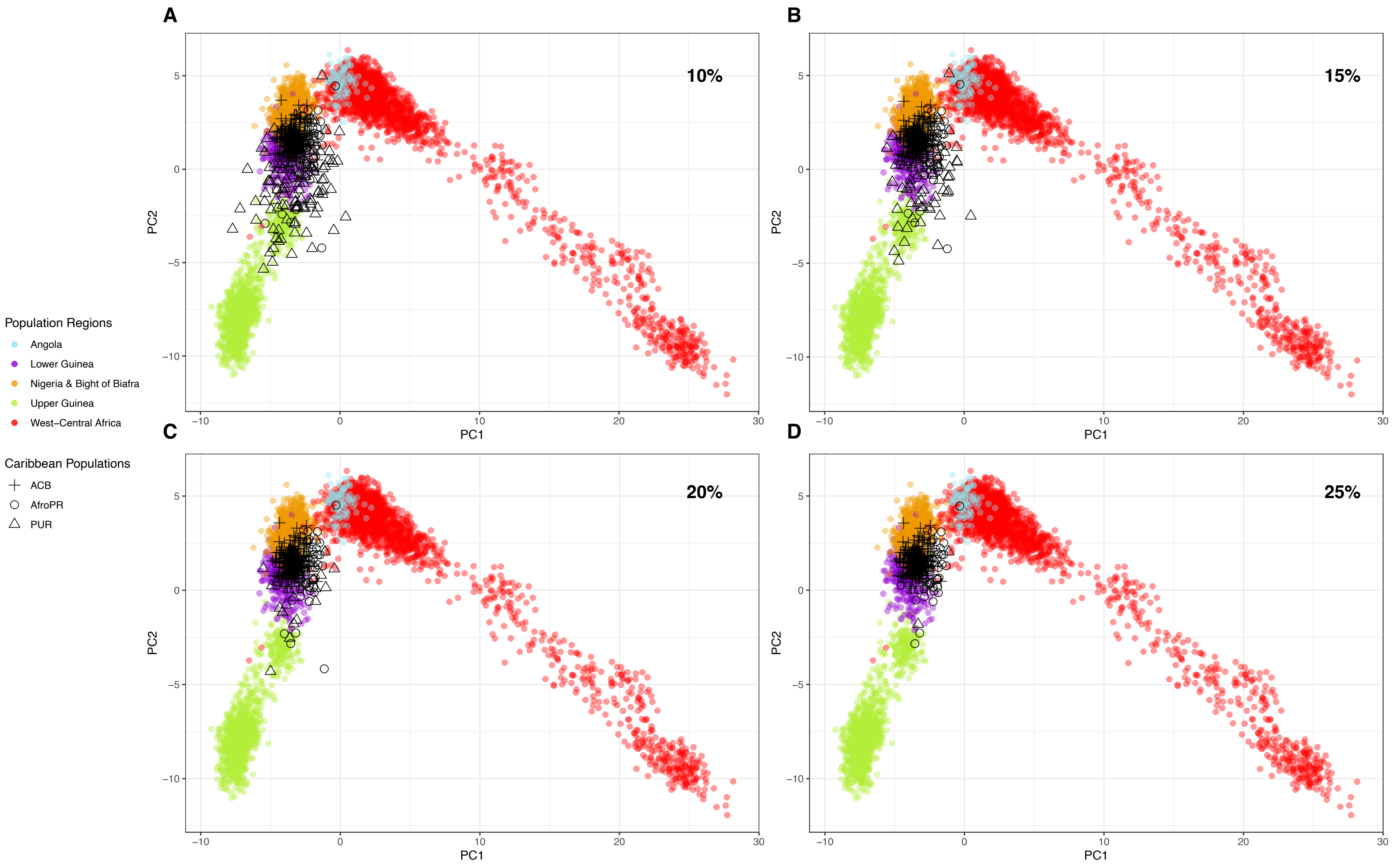


**Figure S7: Subcontinental African ancestry MDS analysis (PC 1 vs. 2) at varying minimum thresholds of African ancestry.** Colored circles represent individual haplotypes from selected subcontinental African reference populations. Open circles are

individual haplotypes from the study population, open triangles from the 1000 Genomes Project PUR population, and crosses from the 1000 Genomes Project ACB population. (A) MDS results for all haplotypes calculated by RFMix to have ≥10% African ancestry. (AfroPR: N=60/60, PUR: N=108/208, ACB: N=192/192) (B) MDS results for all haplotypes calculated by RFMix to have ≥15% African ancestry. (AfroPR: N=58/60, PUR: N=54/208, ACB: N=192/192) (C) MDS results for all haplotypes calculated by RFMix to have ≥20% African ancestry. (AfroPR: N=58/60, PUR: N=22/208, ACB: N=192/192) (D) MDS results for all haplotypes calculated by RFMix to have ≥25% African ancestry. (AfroPR: N=58/60, PUR: N=8/208, ACB: N=192/192)

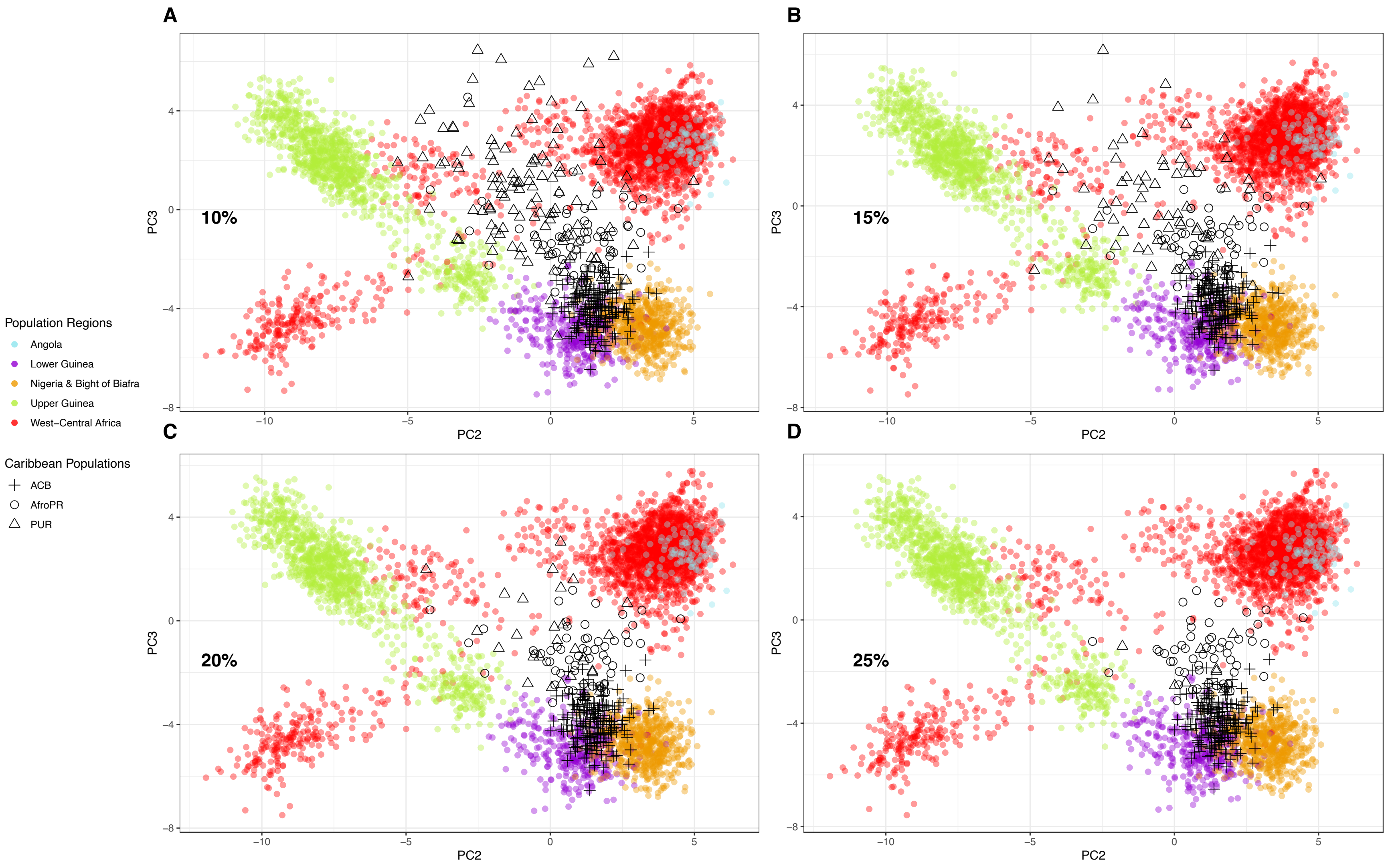


**Figure S8: Subcontinental African ancestry MDS analysis (PC 2 vs. 3) at varying minimum thresholds of African ancestry.** Colored circles represent individual haplotypes from selected subcontinental African reference populations. Open circles are

individual haplotypes from the study population, open triangles from the 1000 Genomes Project PUR population, and crosses from the 1000 Genomes Project ACB population. (A) MDS results for all haplotypes calculated by RFMix to have ≥10% African ancestry. (AfroPR: N=60/60, PUR: N=108/208, ACB: N=192/192) (B) MDS results for all haplotypes calculated by RFMix to have ≥15% African ancestry. (AfroPR: N=58/60, PUR: N=54/208, ACB: N=192/192) (C) MDS results for all haplotypes calculated by RFMix to have ≥20% African ancestry. (AfroPR: N=58/60, PUR: N=22/208, ACB: N=192/192) (D) MDS results for all haplotypes calculated by RFMix to have ≥25% African ancestry. (AfroPR: N=58/60, PUR: N=8/208, ACB: N=192/192)

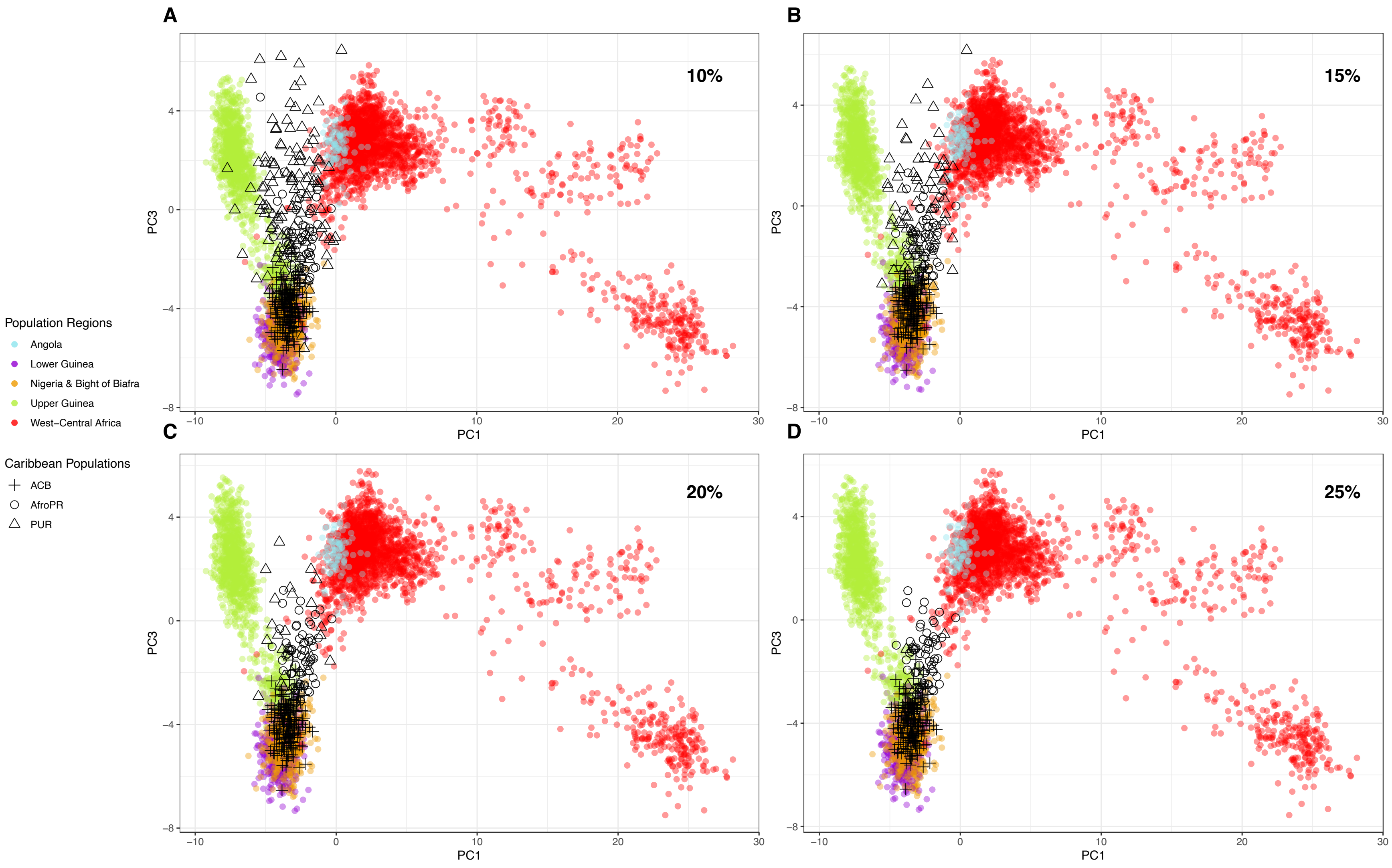


**Figure S9: Subcontinental African ancestry MDS analysis (PC 1 vs. 3) at varying minimum thresholds of African ancestry.** Colored circles represent individual haplotypes from selected subcontinental African reference populations. Open circles are

individual haplotypes from the study population, open triangles from the 1000 Genomes Project PUR population, and crosses from the 1000 Genomes Project ACB population. (A) MDS results for all haplotypes calculated by RFMix to have ≥10% African ancestry. (AfroPR: N=60/60, PUR: N=108/208, ACB: N=192/192) (B) MDS results for all haplotypes calculated by RFMix to have ≥15% African ancestry. (AfroPR: N=58/60, PUR: N=54/208, ACB: N=192/192) (C) MDS results for all haplotypes calculated by RFMix to have ≥20% African ancestry. (AfroPR: N=58/60, PUR: N=22/208, ACB: N=192/192) (D) MDS results for all haplotypes calculated by RFMix to have ≥25% African ancestry. (AfroPR: N=58/60, PUR: N=8/208, ACB: N=192/192)

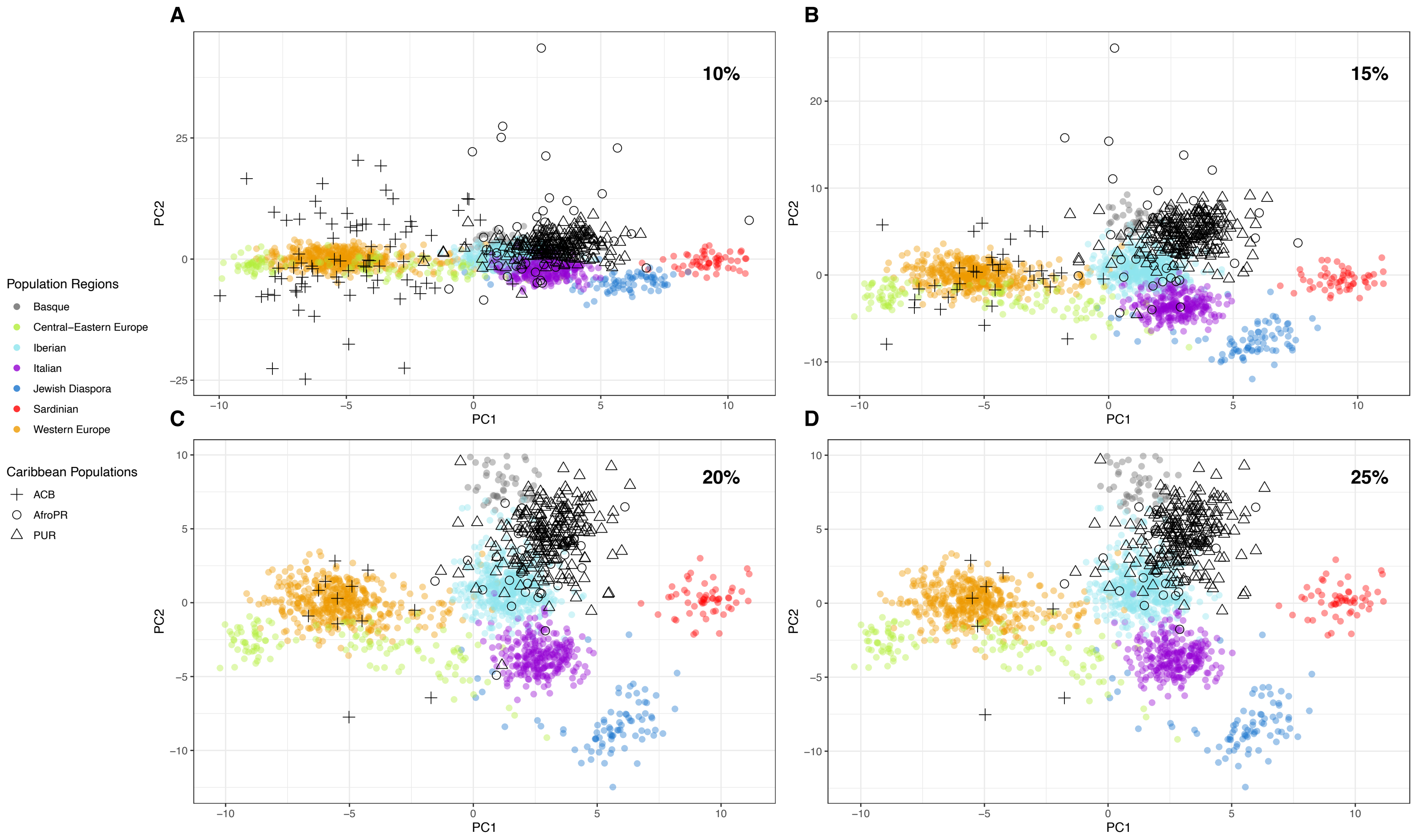


**Figure S10: Subcontinental European ancestry MDS analysis (PC 1 vs. 2) at varying minimum thresholds of European ancestry.** Colored circles represent individual haplotypes from selected subcontinental European reference populations. Open circles are individual haplotypes from the study population, open triangles from the 1000 Genomes Project PUR population, and crosses from

the 1000 Genomes Project ACB population. (A) MDS results for all haplotypes calculated by RFMix to have ≥10% European ancestry. (AfroPR: N=60/60, PUR: N=208/208, ACB: N=76/192) (B) MDS results for all haplotypes calculated by RFMix to have ≥15% European ancestry. (AfroPR: N=48/60, PUR: N=208/208, ACB: N=30/192) (C) MDS results for all haplotypes calculated by RFMix to have ≥20% European ancestry. (AfroPR: N=40/60, PUR: N=208/208, ACB: N=12/192) (D) MDS results for all haplotypes calculated by RFMix to have ≥25% European ancestry. (AfroPR: N=38/60, PUR: N=208/208, ACB: N=6/192)

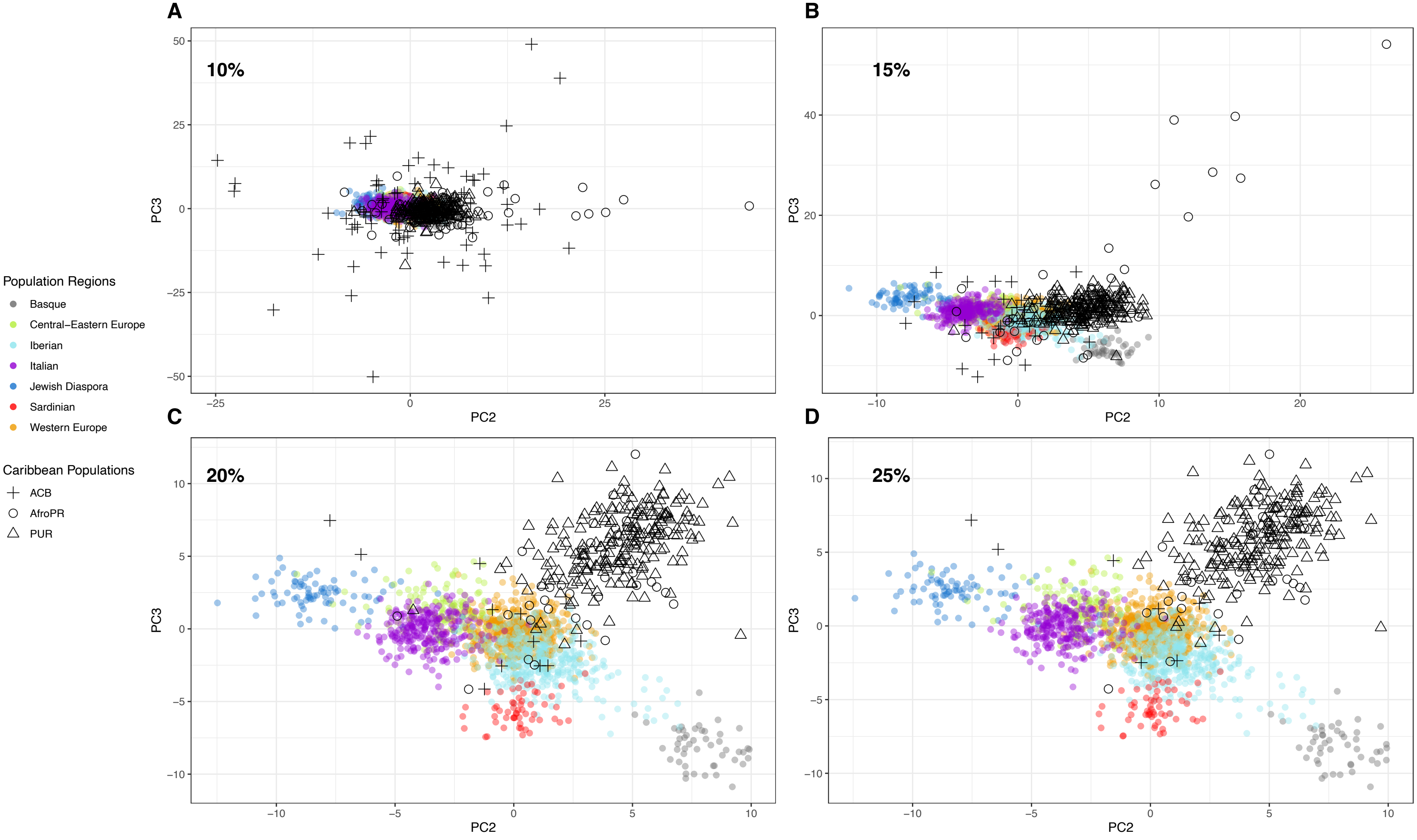


**Figure S11: Subcontinental European ancestry MDS analysis (PC 2 vs. 3) at varying minimum thresholds of European ancestry.** Colored circles represent individual haplotypes from selected subcontinental European reference populations. Open circles are individual haplotypes from the study population, open triangles from the 1000 Genomes Project PUR population, and crosses from

the 1000 Genomes Project ACB population. (A) MDS results for all haplotypes calculated by RFMix to have ≥10% European ancestry. (AfroPR: N=60/60, PUR: N=208/208, ACB: N=76/192) (B) MDS results for all haplotypes calculated by RFMix to have ≥15% European ancestry. (AfroPR: N=48/60, PUR: N=208/208, ACB: N=30/192) (C) MDS results for all haplotypes calculated by RFMix to have ≥20% European ancestry. (AfroPR: N=40/60, PUR: N=208/208, ACB: N=12/192) (D) MDS results for all haplotypes calculated by RFMix to have ≥25% European ancestry. (AfroPR: N=38/60, PUR: N=208/208, ACB: N=6/192)

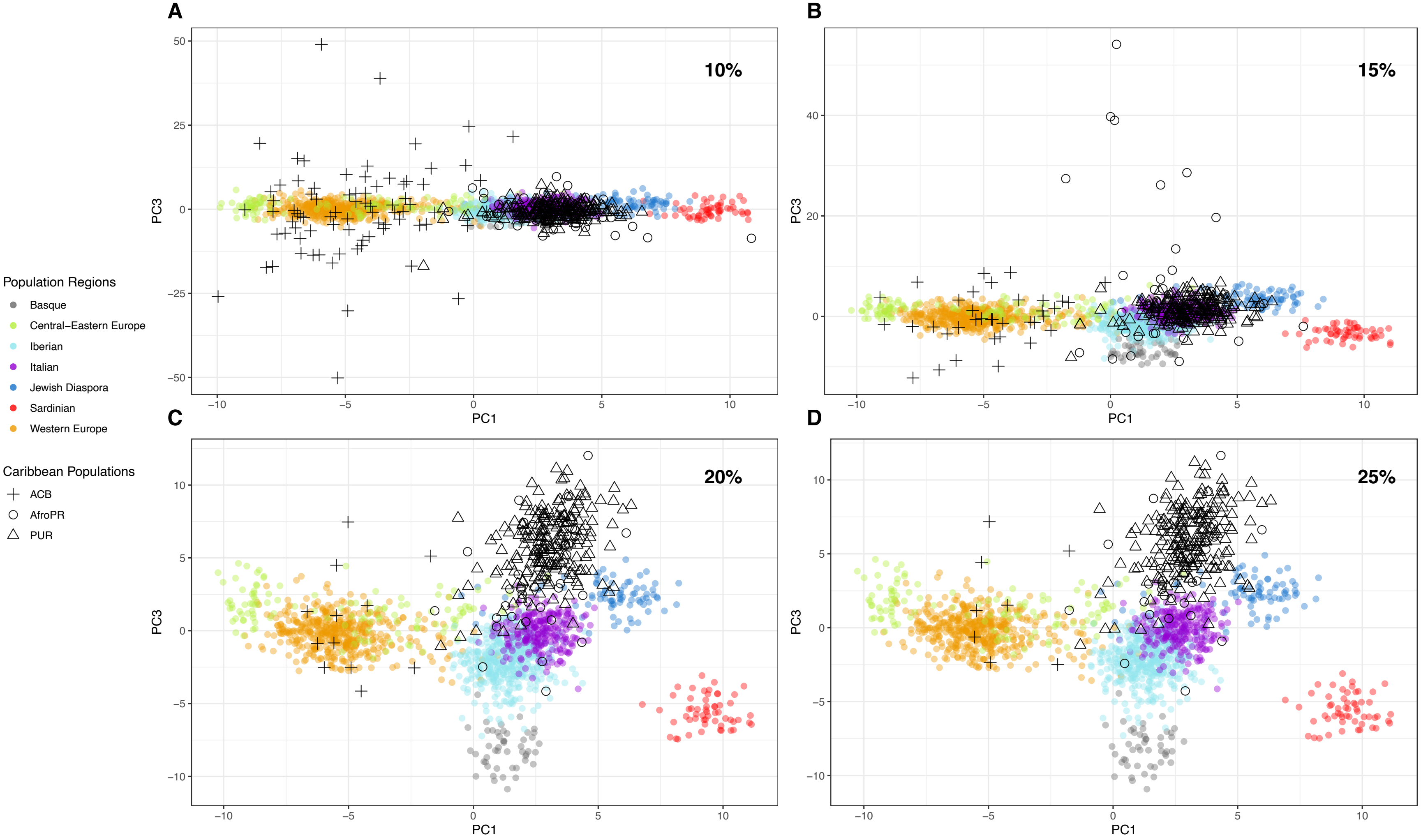


**Figure S12: Subcontinental European ancestry MDS analysis (PC 1 vs. 3) at varying minimum thresholds of European ancestry.** Colored circles represent individual haplotypes from selected subcontinental European reference populations. Open circles are individual haplotypes from the study population, open triangles from the 1000 Genomes Project PUR population, and crosses from

the 1000 Genomes Project ACB population. (A) MDS results for all haplotypes calculated by RFMix to have ≥10% European ancestry. (AfroPR: N=60/60, PUR: N=208/208, ACB: N=76/192) (B) MDS results for all haplotypes calculated by RFMix to have ≥15% European ancestry. (AfroPR: N=48/60, PUR: N=208/208, ACB: N=30/192) (C) MDS results for all haplotypes calculated by RFMix to have ≥20% European ancestry. (AfroPR: N=40/60, PUR: N=208/208, ACB: N=12/192) (D) MDS results for all haplotypes calculated by RFMix to have ≥25% European ancestry. (AfroPR: N=38/60, PUR: N=208/208, ACB: N=6/192)
